# Coupling of Ca^2+^-triggered unclamping and membrane fusion during neurotransmitter release

**DOI:** 10.1101/2021.06.16.448753

**Authors:** Zachary A. McDargh, Anirban Polley, Jin Zeng, Ben O’Shaughnessy

## Abstract

Neurotransmitter (NT) release is accomplished by a machinery that unclamps fusion in response to calcium and then fuses the synaptic vesicle and plasma membranes. These are often thought of as distinct tasks assigned to non-overlapping components. Vesicle release rates have a power law dependence on [Ca^2+^] with an exponent of 3-5, long taken to indicate that 3-5 Ca^2+^ ions bind the calcium sensor Synaptotagmin to trigger release. However, dependencies at low [Ca] are inconsistent with simple sequential binding to a single Ca^2+^ sensor followed by a final fusion step. Here we developed coarse-grained molecular dynamics simulations of the NT release machinery accounting for Synaptotagmin-mediated unclamping and SNARE-mediated fusion. Calcium-triggered unclamping and SNARE-mediated fusion emerged from simulations as contemporaneous, coupled processes. Increasing cytosolic [Ca^2+^], the instantaneous fusion rate increased as SNAREpins were progressively and reversibly released by dissociation of Synaptotagmin-SNAREpin complexes. Simulations reproduced the observed dependence of release rates on [Ca^2+^], but the power law was unrelated to the number of Ca^2+^ ions required. Action potential-evoked vesicle release probabilities depended on the number of transiently unclamped SNAREpins, explaining experimental dependencies of release probabilities on both unclamping and membrane-fusing machinery components. These results describe a highly cooperative NT release machinery with intrinsically inseparable unclamping and membrane-fusing functionalities.

## Introduction

Neurotransmission relies on secretion of neurotransmitters (NTs) at synapses, accomplished by a specialized machinery that responds to action potential-evoked calcium influx at the presynaptic terminal. On submillisecond timescales, the machinery releases SNARE proteins to fuse synaptic vesicle and plasma membranes and release NTs (Brunger et al., 2018a; Sudhof, 2013) into the synaptic cleft that bind post-synaptic cell receptors (Ehlers et al., 1996).

The kinetics of synaptic transmission have been characterized by electrophysiological techniques. Following initiation of a presynaptic action potential (AP), excitatory postsynaptic currents (EPSCs) are measured due to simultaneous release from multiple synaptic contacts between two neurons (Branco and Staras, 2009), or at individual synapses (Augustine and Charlton, 1986; Borst and Sakmann, 1998; Kawaguchi and Sakaba, 2017). Measurement of the quantal release per vesicle from spontaneous release events (Bollmann et al., 2000; Schneggenburger and Neher, 2000) and the size of the readily-releasable pool (RRP) (Rosenmund and Stevens, 1996) enables conversion of EPSCs to vesicle release rates (Bollmann et al., 2000; Schneggenburger and Neher, 2000) and computation of the vesicle release probability *P*_ves_, the fraction of RRP vesicles released following an AP.

These studies showed that the AP signal of 0.5-2.0 ms duration activates a pre-synaptic calcium transient typically lasting ~ 0.2 – 1 ms (Borst and Sakmann, 1998; Dittman and Ryan, 2019; Neher and Sakaba, 2008) that elicits a post-synaptic response with a ~ 0.5-2 ms delay measured from the AP peak to the start of the EPSC (Katz and Miledi, 1965; Sabatini and Regehr, 1996). Decades ago Katz and colleagues attributed the delay at the frog neuromuscular junction primarily to NT release (Katz and Miledi, 1965). Another key finding is that *P*_ves_ is usually small (Branco and Staras, 2009; Dittman and Ryan, 2019), attributed to effects such as [Ca^2+^] levels at the release site (Bohme et al., 2018; Dittman and Ryan, 2019; Fioravante and Regehr, 2011) and vesicle priming factors (Korber and Kuner, 2016; Rosenmund et al., 2002).

The NT release machinery is more directly interrogated by methods that control [Ca^2+^] at the presynaptic terminal, by control of extracellular [Ca^2+^] or flash photolysis to uncage intracellular [Ca^2+^]. Synaptotagmin 1 (Syt) was identified as the Ca^2+^ sensor for synchronous NT release, as mutations altering the Ca^2+^ binding affinity of Syt proportionally altered the Ca^2+^ sensitivity of release (Fernandez-Chacon et al., 2001; Rhee et al., 2005). At many synapses, EPSC amplitude increases with a power law dependence on [Ca^2+^] with an exponent of 3-5, (Augustine and Charlton, 1986; Neher and Sakaba, 2008; Rahamimoff and Dodge, 1969), widely interpreted as signifying that ~3-5 Ca^2+^ ions cooperatively trigger release (Augustine and Charlton, 1986; Rahamimoff and Dodge, 1969). At large [Ca^2+^] the EPSC amplitude shows a plateau, but its origin is not established (Acuna et al., 2014; Sakaba et al., 2005; Wang et al., 2008).

A major challenge is to understand how the NT release machinery leads to these behaviors. Many components are now identified. Prior to Ca^2+^ entry Syt is thought to clamp fusion by the neuronal SNARE proteins (Sollner et al., 1993; Weber et al., 1998) and to synchronize release with the Ca^2+^ stimulus (Nishiki and Augustine, 2004a, b). Other components include Munc18 and Munc13, which regulate SNARE complex assembly (Lai et al., 2017; Ma et al., 2015) and Complexin, with reported clamping and facilitating roles (Giraudo et al., 2006; Ramakrishnan et al., 2020; Xue et al., 2010). While phenomenological models were developed (Bollmann et al., 2000; Lou et al., 2005; Schneggenburger and Neher, 2000; Sun et al., 2007), molecularly detailed quantitative models are not available. One model is that Syt-SNARE interactions inhibit SNARE complexation (Grushin et al., 2019), until Ca^2+^ binding to Syt releases the SNAREs for fusion. Another proposal is that Syt clamps fusion by chaining SNARE complexes together via opposing SNARE-binding interfaces on the two Ca^2+^-binding C2 domains of Syt (Brunger et al., 2018b; Zhou et al., 2015).

A third proposal stems from the finding that Syt oligomerizes into ~30 nm rings on anionic lipid monolayers that spontaneously disassemble at physiological [Ca^2+^] (Wang et al., 2014; Wang et al., 2017; Zanetti et al., 2016), suggesting Syt rings could clamp fusion by spacing the vesicle and plasma membranes until Ca^2+^ triggers ring disassembly and fusion. Mutations disabling Syt oligomerization disrupted clamping in PC12 cells and cortical neurons (Bello et al., 2018; Tagliatti et al., 2020) and *in vitro* (Ramakrishnan et al., 2018; Ramakrishnan et al., 2020), and abolished a symmetric arrangement at the vesicle-plasma membrane interface (Li et al., 2019b). However, recent EPR studies found no evidence of Syt oligomerization (Nyenhuis et al., 2019).

Following Ca^2+^-mediated unclamping the unfettered SNAREs mediate fusion, but the mechanism is controversial. Fusion is commonly thought driven by the SNARE complex zippering energy (Gao et al., 2012; Ma et al., 2015), but SNARE linker domains (LDs) may be flexible (Kim et al., 2002; Lakomek et al., 2019) and incapable of storing bending energy to press the membranes together. Our previous simulations suggest an entirely different picture, in which fully zippered SNARE complexes drive fusion through entropic forces that push SNARE complexes outwards and pull the membranes together (McDargh et al., 2018; Mostafavi et al., 2017).

Here we develop a molecularly detailed mathematical model of the NT release machinery that presents a unified account of Ca^2+^-triggered unclamping and SNARE-mediated membrane fusion. The model incorporates a core machinery consisting of a Syt ring at the vesicle-plasma membrane interface bound by SNARE complexes according to the crystal structure (Zhou et al., 2015). Components with unknown architecture are omitted, but many of our qualitative conclusions are independent of the detailed architecture. Ca^2+^ influx, ring disassembly, unclamping and fusion are tracked. We find the synaptic delay and the high [Ca^2+^] release rate plateau are dominated by the unclamping and fusion times, respectively. The model reproduces the power law dependence of release rates on [Ca^2+^], but shows this dependence is unrelated to the number of [Ca^2+^] ions required for NT release. Consistent with experiment (Acuna et al., 2014; Arancillo et al., 2013; Ruiter et al., 2019) we find the release probability *P*_ves_ depends on both unclamping and fusion, since unclamping provides a brief window when fusion may occur before reclamping. Thus, NT release is accomplished by a highly cooperative machinery whose Ca^2+^-triggered unclamping and membrane-fusing functionalities are intrinsically inseparable.

## Results

### Model

We developed coarse-grained molecular dynamics simulations of Ca^2+^-triggered fusion of synaptic vesicles with the presynaptic plasma membrane (PM). The simulations represent Syt and neuronal SNARE complexes (SNAREpins), Fig. 1A. Simulated vesicles host 20 Syt molecules, comparable to the ~15-20 reported in synaptic vesicles (Takamori et al., 2006; Wilhelm et al., 2014), initially assembled into a ring, Figs. 1B, 2A. Ten trans-SNARE complexes are bound to the Syt ring via the primary interface from the Syt-SNARE complex crystal structure (Zhou et al., 2015), the maximum without steric clashes. This may maximize the fusion rate following Ca^2+^ entry, since simulations suggested SNARE-mediated fusion rates increase with more SNAREs (McDargh et al., 2018; Mostafavi et al., 2017).

**Figure 1:**
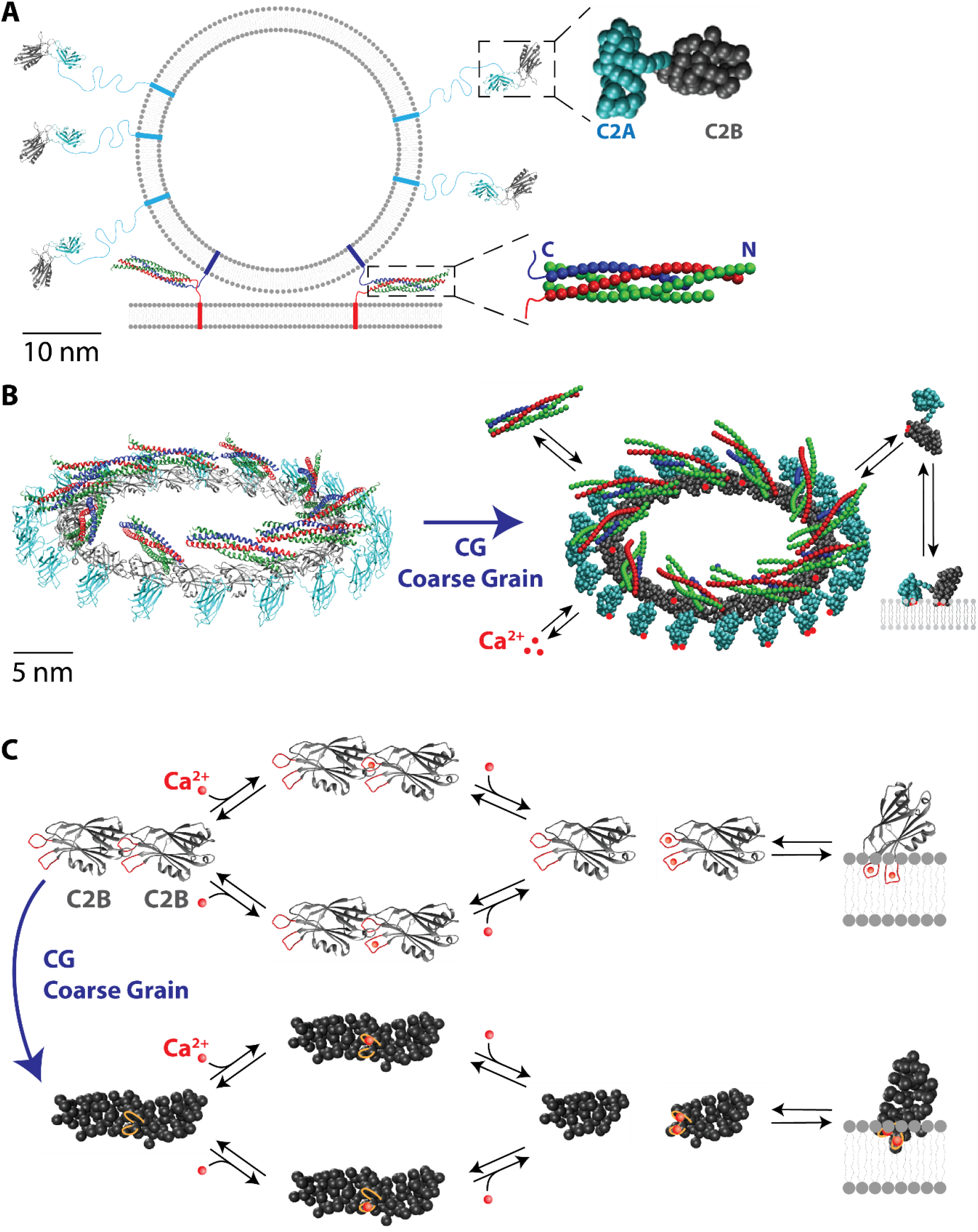
Coarse-grained model of a minimal neurotransmitter release machinery. **(A)** Left: Schematic of a vesicle docked to the PM via trans-SNARE complexes, to scale. The SNARE complex comprises VAMP (blue), Syntaxin (red), and SNAP-25 (green). Right: CG models of Syt and the SNARE complex. **(B)** Left: Syt ring reconstructed from electron micrographs (Wang et al., 2014) with SNARE complexes (PDB ID: 3HD7 (Stein et al., 2009)) docked to every other Syt molecule. Right: CG representation. The model incorporates Syt-SNARE, Syt-Ca^2+^, Syt-ring and Syt-PM association/dissociation kinetics. **(C)** When both C2B domain Ca^2+-^binding sites are occupied, Syt neighbors in the ring dissociate and the C2B domain can bury its Ca^2+^-binding loops into the plasma membrane. C2A domain omitted for clarity.

To access the timescales of NT release, we use highly coarse-grained (CG) representations (see Supplementary Material). The simulated SNARE complex comprises one helix from the vesicle-associated VAMP, two from SNAP-25, and one from the PM-associated Syntaxin (Stein et al., 2009) while Syt includes the C2A and C2B domains, connected to the vesicle-associated transmembrane domain (TMD) by a 60-residue linker domain (LD), Fig. 1A. One bead represents four residues in alpha helices and beta sheets, and two residues in unstructured loops, Fig. 1A (McDargh et al., 2018; Mostafavi et al., 2017). The LD of Syt, and of Syntaxin and VAMP including the unassembled (unzippered) portions, are assumed unstructured (Kim et al., 2002; Lakomek et al., 2019), represented by worm-like chains with parameters depending on the degree of unzippering. The Syt C2AB domain and the assembled part of the SNARE complex are undeformable. The 40 nm diameter vesicle and planar membranes are continuous non-deformable surfaces. We used simulations to compute the instant of membrane fusion and release, defined as the time when the membrane interaction energy *E*_mb_ first exceeds the fusion barrier, *E*_fusion_ = 20 *kT* (Francois-Martin et al., 2017).

We implemented dynamic SNARE complex zippering/unzipping, binding/dissociation of Ca^2+^ ions to the Syt C2A and C2B domains (Radhakrishnan et al., 2009), Syt-Syt binding/dissociation (Wang et al., 2014), burying/unburying of Syt C2A and C2B domain Ca^2+^-binding loops into the PM (Ma et al., 2017; Perez-Lara et al., 2016), and Syt-SNARE binding/dissociation (Zhou et al., 2015), Fig. 1B.

Syt has three Ca^2+^-binding sites on the C2A domain (Ubach et al., 1998), and two on the C2B domain (Fernandez et al., 2001). Ca^2+^ cooperatively triggers Syt-liposome binding (Radhakrishnan et al., 2009) and Syt rings spontaneously disassemble at physiological [Ca^2+^] (Wang et al., 2014), Fig. 1C. The model assumes insertion of the C2A or C2B Ca^2+^-binding loops into the PM requires all Ca^2+^-binding sites to be bound, Syt-Syt bonds break instantly when both C2B Ca^2+^-binding sites are occupied, and re-binding requires at least one unoccupied Ca^2+^-binding site, Fig. 1C.

Syt-SNARE binding at the primary interface and Syt-Syt oligomerization in the absence of Ca^2+^ occur within a capture distance, with dissociation rates that reproduce the observed respective dissociation constants (Wang et al., 2017; Zhou et al., 2017), Figs. 1B, C and S2.

SNAREpins assemble and disassemble layer by layer at rates *k*_zip_ = *k*_0_ exp[−Δ*E*_zip_/kT] and *k*_unzip_ = *k*_0_ = 10^6^ s^−1^ (Kubelka et al., 2004), where the zippering energy Δ*E*_zip_ is known from the measured SNARE zippering free energy landscape (Gao et al., 2012; Ma et al., 2015) and the stretching energies of the uncomplexed LDs. Uncomplexed beads are assumed unstructured. Disassembly in the C-terminal domain (layers +5 to +8) removes a bead from all four SNARE helices, consistent with observed C-terminal fraying (Ma et al., 2015).

For model details and simulation parameters, see Supplementary Material and Tables S1-S2.

### Synaptotagmin rings clamp fusion by spacing membranes and blocking SNAREpin reorganization

Simulations supported the hypothesis that oligomeric Syt rings clamp fusion before Ca^2+^ entry into axon terminals. With [Ca^2+^]=0.1 μM, a typical presynaptic basal value in neurons (Ermolyuk et al., 2013; Jackson and Redman, 2003), all Syt rings remained intact with unbroken Syt-Syt bonds and and no fusion occurred (100 simulations, total 200 ms simulation time), Fig. 2A. Those Syt monomers that spontaneously dissociated re-associated within ~1 μs.

**Figure 2:**
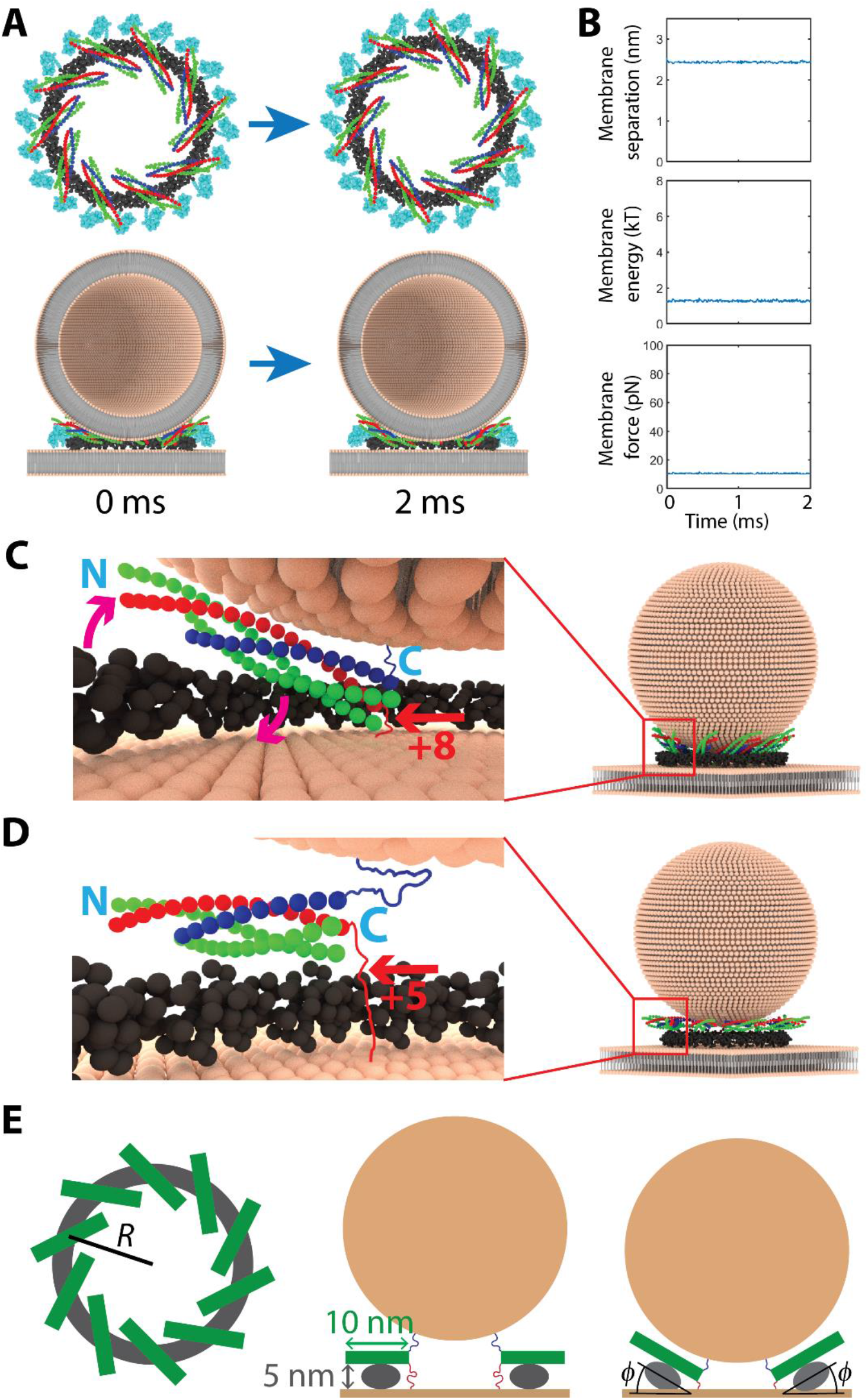
Syt rings clamp fusion by spacing membranes and blocking SNAREpin reorganization. **(A)** Top views of the Syt ring with bound SNAREpins (top) and side views with a vesicle cross-section (bottom) during a typical simulation with [Ca^2+^] = 0.1 μM. Over 100 runs, each lasting 2 ms, no fusion occurred. Membranes are depicted schematically, with explicit lipids. **(B)** Membrane separation at point of closest approach, membrane energy and membrane force vs. time for the simulation of (A). Vertical scales chosen for comparison with Fig. 4C. **(C)** Snapshot of a simulated vesicle in the clamped configuration. The SNARE motifs fully zipper to layer +8, pulling down the C-terminal end of the SNARE complex. Since the SNARE complex is bound to the Syt ring, the soft ring is twisted (arrows). In (**C**-**E**), Syt C2A domains and linkers are omitted for clarity. **(D)** In simulations with artificially rigid Syt rings, ring twisting is no longer possible. SNAREpins are then maintained in a ‘horizontal’ orientation with C termini a large distance from the plasma membrane. Tension in the Syntaxin LD prevented zippering beyond layer +5. **(E)** Elastic model of the Syt-SNARE ring (SNARE complexes green, Syt grey). Left: top view of ring. Center: unzippered SNAREpins are roughly parallel to the membrane, with raised C-termini. Right: zippering lowers the C-termini, tilting the SNAREs and twisting the Syt ring by angle *ϕ*. This generates elastic stress, as the Syt ring prefers to curve along the long axis of its elliptical cross-section only.

Fusion was clamped by two mechanisms, Fig. 2. First, the Syt ring was a spacer, imposing a membrane separation of at least ~2.5 nm at which the membrane interaction energy *E*_mb_ (1.5 ± 0.1 *kT*) was far from the fusion threshold and the vesicle exerted a small force 12 ± 0.5 pN of electrostatic origin on the PM, well below the ~45 pN in prior simulations lacking Syt (McDargh et al., 2018; Mostafavi et al., 2017).

Second, the Syt ring inhibited fusion by binding the SNAREpins and fixing their location, preventing their spatial reorganization which would otherwise catalyze fusion (see later sections). On average, all 10 SNAREpins remained bound to their associated Syt, with small fluctuations when SNAREs transiently dissociated.

### Due to their flexibility Syt rings permit full SNAREpin zippering in the clamped state

It was recently suggested that the Syt ring could lock SNAREpins into a partially assembled intermediate state that could block fusion, by lifting the SNAREpin C-terminal ends 5 nm from the PM (Grushin et al., 2019). Assembly of the SNARE complexes beyond layer +5 would be prevented, as this would over-stretch the Syntaxin and SNAP-25 LDs. Thus, binding to the Syt ring would lock the SNAREs into this partially unzippered state, Fig. 2D.

Simulations did not support this picture. At resting [Ca^2+^] the SNARE motifs fully zippered to layer +8 by twisting the Syt ring so the SNARE complex C-termini were tilted down towards the PM, preventing the Syntaxin LD from becoming overstretched. Thus, Syt ring flexibility allowed complete zippering, with only the juxtamembrane LDs remaining unstructured, Fig. 2C. Zippering was only restrained when we endowed Syt rings with artificial rigidity: then SNAREs indeed could not zipper beyond layer +5 as predicted, Fig. 2D (Grushin et al., 2019).

This underlines the importance of Syt ring flexibility. Measurements of Syt rings in solution revealed a broad size distribution (Wang et al., 2017; Zhu et al., 2021) which is directly related to the ring stiffness or persistence length *l*_*p*_ (Zhu et al., 2021) and implies *l*_*p*_ ~40 - 170 nm (Wang et al., 2017; Zhu et al., 2021). We used a representative value, *lp* = 70 nm (Table S1), so Syt ring shapes fluctuated considerably, allowing complete SNARE complex assembly, Fig. 2C.

We quantified these effects using a simple continuum elastic model of the Syt ring (see Supplementary Material). Its starting point is the bending energy of a Syt-SNARE ring of radius *R* with SNAREs tilted by angle *ϕ*, Fig. 2E, subject to material curvatures *c*_1_ = cos(*ϕ*) / *R* and *c*_2_ = sin(*ϕ*) /*R* parallel and perpendicular to the membrane plane in the unstressed configuration, respectively:

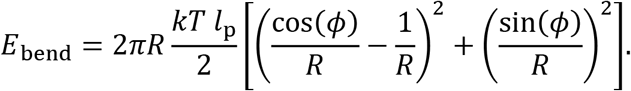

Here the ring has spontaneous curvature 1/*R* in the material direction parallel to the membrane in the un-twisted state, and zero spontaneous curvature in the out-of-plane direction. Given the ~10 nm SNAREpin length and ~5 nm C2B domain thickness, a tilt *ϕ* ≈ 45° would lower the SNAREpin C-terminal to within ~2 nm of the PM, allowing full zippering, Fig 2E. From the expression above, this costs bending energy *E*_bend_ ≈ 10 *kT*, far less than the ~40 *kT* to unzipper even one layer of the 10 SNAREpins.

Thus, Syt rings are far too soft to sustain significant SNARE complex unzippering in the initial clamped state. Note this does not compromise the ability of the Syt ring to clamp fusion, Fig. 2.

### Ca^2+^ entry disassembles Syt rings and releases SNAREpins by dissociating Syt-SNARE complexes

To examine the molecular mechanism of Ca^2+^-triggered unclamping, we simulated Ca^2+^ uncaging in the calyx of Held, when [Ca^2+^] is abruptly elevated throughout the axon terminal. This method was used at the calyx of Held, Schaffer collaterals and other synapses (Burgalossi et al., 2010; Sakaba, 2008; Schneggenburger and Neher, 2000).

We ran 100 ~ 10 ms simulations per [Ca^2+^] value in the range 5-40 μM. Following uncaging, Ca^2+^ binding to Syt C2B domains triggered Syt-Syt dissociation, progressively disassembling the Syt ring that had prevented vesicle-plasma membrane contact, Figs. 3A, B. A second inhibition removed was spatial confinement of the SNARE complexes: Ca^2+^ binding dissociated Syt-SNARE complexes, unclamping the SNAREs. We defined a SNAREpin to be clamped if it was bound to a Syt monomer that itself was bound to at least one other Syt monomer. A vesicle was defined to be clamped if four or more of its associated SNAREpins were clamped. The number of unclamped SNAREpins increased with time, Fig. 3B, so that vesicles became unclamped after 850±370 μs for [Ca^2+^]=25 μM (n=100 simulations). At lower [Ca^2+^], Ca^2+^ bound Syt more slowly so unclamping was slower, requiring 6.6 ± 1.1 ms (n=100) at [Ca^2+^]=5 μM.

**Figure 3:**
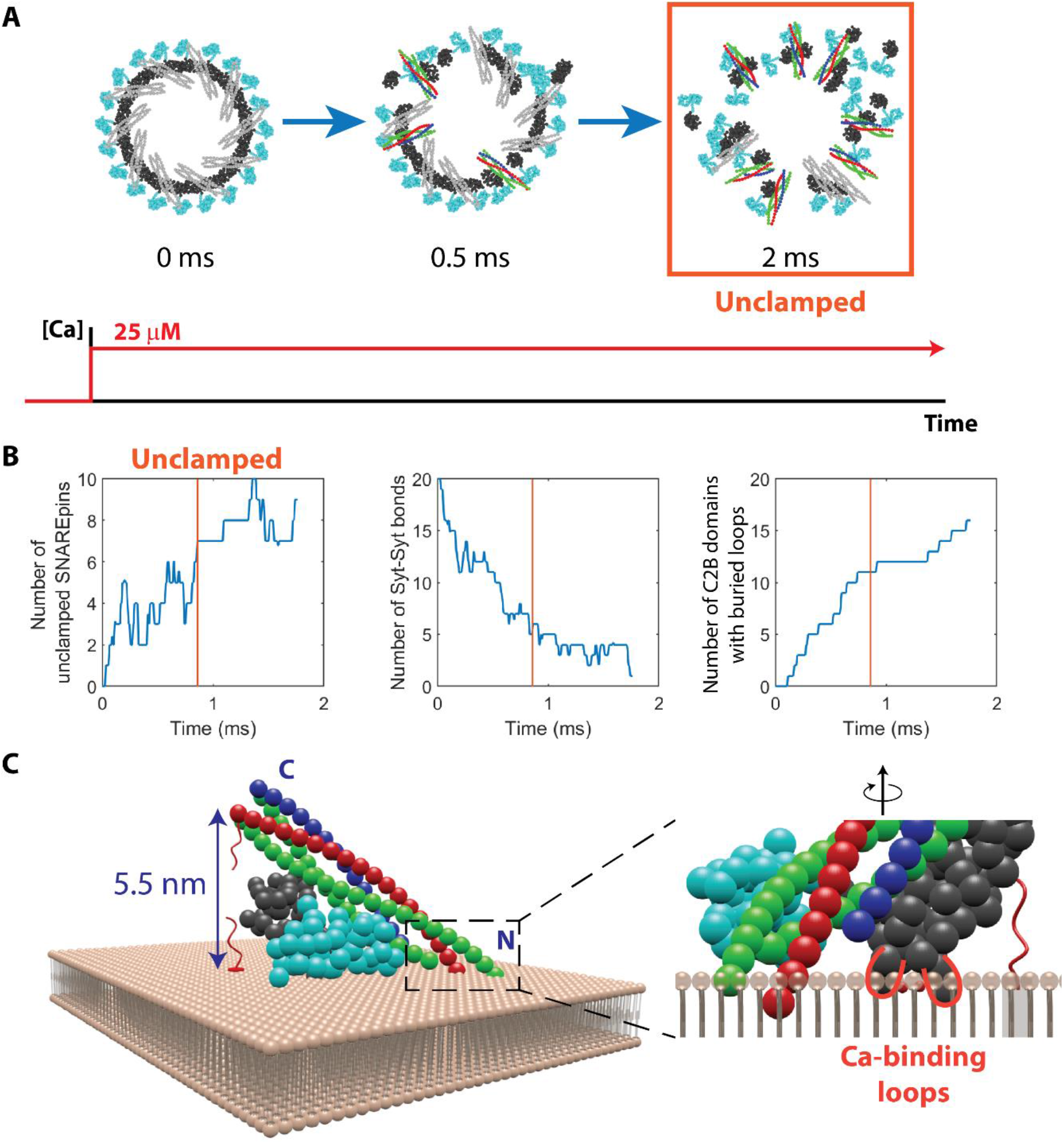
Calcium entry triggers unclamping by disassembly of the Syt ring and dissociation of Syt-SNARE complexes. (A) Snapshots from a typical Ca^2+^ uncaging simulation with [Ca^2+^]= 25 μM. Ca^2+^ binding progressively dissociated Syt-Syt bonds, disassembled the ring, and released the SNAREpins. Clamped SNAREpins shown gray. The vesicle became unclamped after 2 ms. (B) Typical time courses of the numbers of unclamped SNAREpins, Syt-Syt bonds, and Syt C2B domains whose Ca^2+^-binding loops are buried in the plasma membrane ([Ca^2+^] = 25 μM). Orange line: unclamping time. (C) Prohibited configuration of the Syt-SNARE complex, with the C2B Ca^2+^-binding loops (red, see blow-up) inserted into the PM. The Syt C2B domain is in the experimentally observed Ca^2+^-dependent membrane-bound state (Perez-Lara et al., 2016). This configuration would over-stretch the Syntaxin LD and was not seen in simulations. Instead, Ca^2+^ entry triggered SNAREpin dissociation from C2B. This hypothetical configuration would also provoke a steric clash at the N-terminus (right).

Syt-SNAREpin dissociation was driven by Ca^2+^-dependent insertion of the C2B Ca^2+^-binding loops into the PM, which could occur only if the C2B was no longer bound to its SNAREpin partner. This is seen from structures of the Syt-SNARE complex (Zhou et al., 2015) and of membrane-bound C2B domains reconstructed from EPR recordings (Perez-Lara et al., 2016). Loop insertion into the PM rotates the C2 domain so the C-terminus of a bound SNAREpin tilts away from the membrane, Fig. 3C. This tilting cannot be accomplished with fully zippered SNARE motifs, as it would overstretch the Syntaxin LD. Thus, either unzippering occurs, or the Ca^2+^-binding loops are prevented from penetrating the PM. Since zippering and Ca^2+^-dependent Syt-PM binding are highly energetically favorable (~4 kT per layer and ~10 kT, respectively (Ma et al., 2017; Ma et al., 2015)) whereas the Syt-SNARE primary interface is relatively weak (~4.8 kT (Zhou et al., 2017)), the SNARE complex will instead dissociate from its Syt partner.

Insertion of the loops into the PM prevented re-assembly of the Syt ring, since the Syt oligomerization interface is located on the same surface of the C2B domain as the Ca^2+^-binding loops, and insertion was effectively irreversible on the simulation timescales (*k*_off_~1 s^−1^, Table S1).

### Entropic forces organize SNARE complexes into a ring and drive membrane fusion

Unclamped SNAREpins spontaneously assembled into a ring-like organization with radially oriented SNAREpins. Initially, when bound to the Syt ring, the mean orientation angle was ⟨*θ*⟩ = 52 ± 0.6°, imposed by the Syt-SNARE complex structure, Fig. 4A. SNAREpins progressively dissociated, and in steady state became radially oriented with ⟨*θ*⟩ = 0. ±8°, Fig. 4B and S4. The time at which ring assembly occurred (relative to the instant of [Ca^2+^] increase) was 6.6 ± 1.1 ms and 1.1 ± 0.5 ms at 5 μM and 25 μM, respectively (*n* = 100 runs) (Supplementary Material). Typically, assembly occurred ~0.2 ms after unclamping.

**Figure 4:**
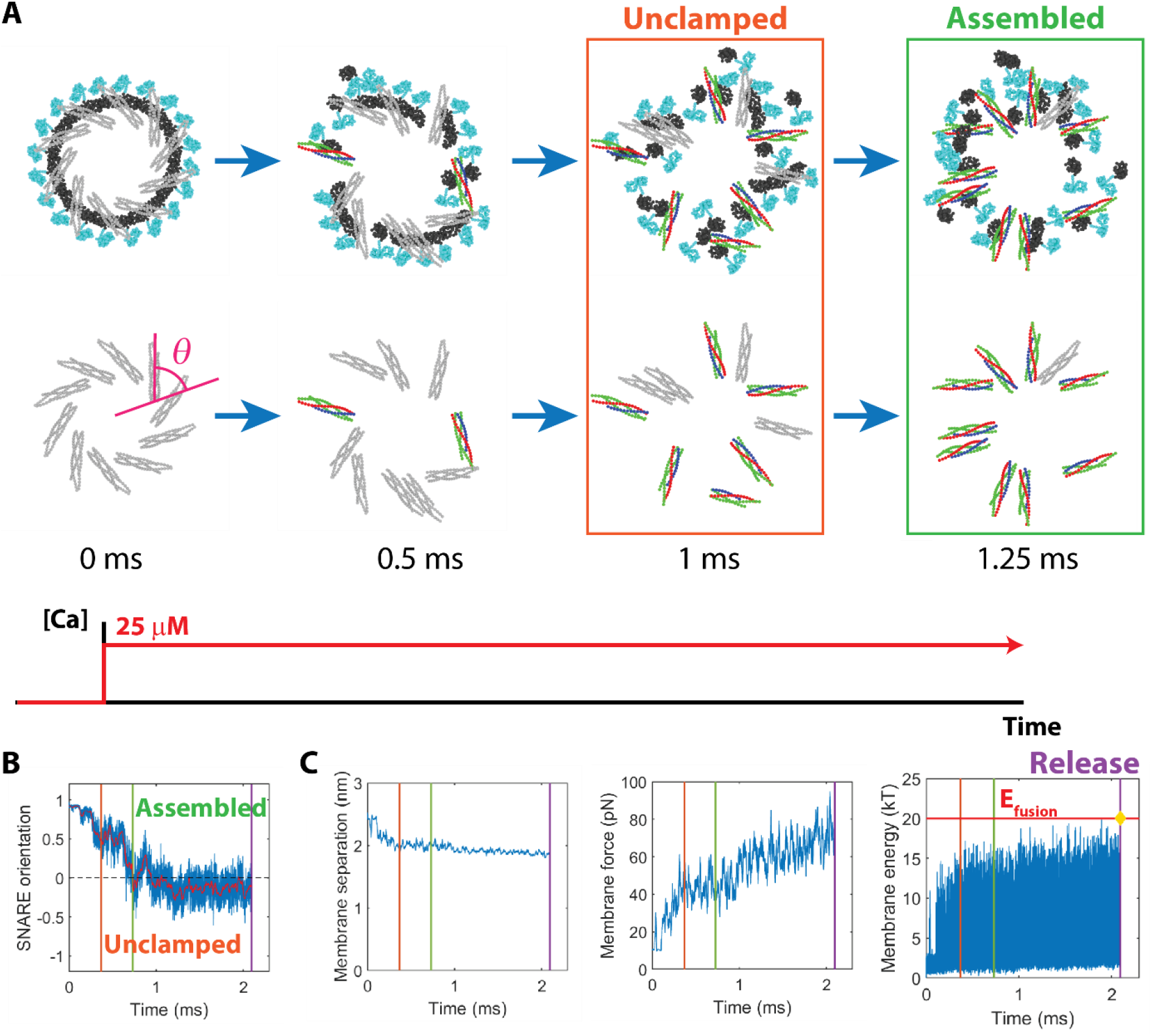
Unclamped SNAREpins self-assemble into a ring that entropically drives membrane fusion. (A) Snapshots from a typical Ca^2+^ uncaging simulation, [Ca^2+^]=25 μM. Following dissociation from the Syt ring and vesicle unclamping, the SNAREpins assembled into a ring after 1.25 ms. Bottom row repeats top row but Syt is omitted. Clamped and unclamped SNAREpins shown gray and in color, respectively. (B) Time course of SNARE orientation angle *θ* (see (A)) averaged over all SNAREs in a typical Ca^2+^ uncaging simulation, [Ca^2+^]=25 μM (blue curve). Red curve: 10 μs moving average. Vesicle unclamping, SNAREpin ring assembly and fusion (purple line) occurred after ~0.4 ms, ~0.7 ms and ~2.1 ms, respectively. (C) Membrane separation, force pressing membranes together and membrane energy for the simulation of (B). Fusion occurred when the energy reached the fusion threshold (yellow diamond).

The forces driving ring assembly were entropic, due to steric interactions among the SNAREpins and membranes. These forces cleared the fusion site of SNAREpins, pushing them outwards and radially aligning them. Organized into an expanded ring, the entropy was increased as the SNAREpins had more freedom to orient laterally and to tilt vertically. Due to vesicle curvature, the ring expansion pulled the vesicle and PM together (McDargh et al., 2018; Mostafavi et al., 2017). As more SNAREpins became unclamped and the SNARE ring progressively assembled, the force pushing the membranes together increased dramatically as did the membrane energy, Fig. 4C. After a long waiting time with the SNAREpin ring in steady state, a fluctuation drove the membrane energy above the fusion barrier, *E*_fusion_ = 20 *kT*.

### Increasing [Ca^2+^] shortens the synaptic delay time in Calyx of Held simulations

The model reproduced the sub-ms delay times measured at the calyx of Held (Bollmann et al., 2000; Lou et al., 2005). We measured the release time (the instant of fusion, Fig. 4C) in multiple Ca^2+^ uncaging simulations, giving a distribution of vesicle release times. Convolving with the average miniature EPSC (mEPSC) measured at the calyx of Held yielded the EPSC, Fig. 5A (Schneggenburger and Neher, 2000). The delay time to the start of the EPSC was measured as the time when 5% of simulated vesicles had released.

**Figure 5:**
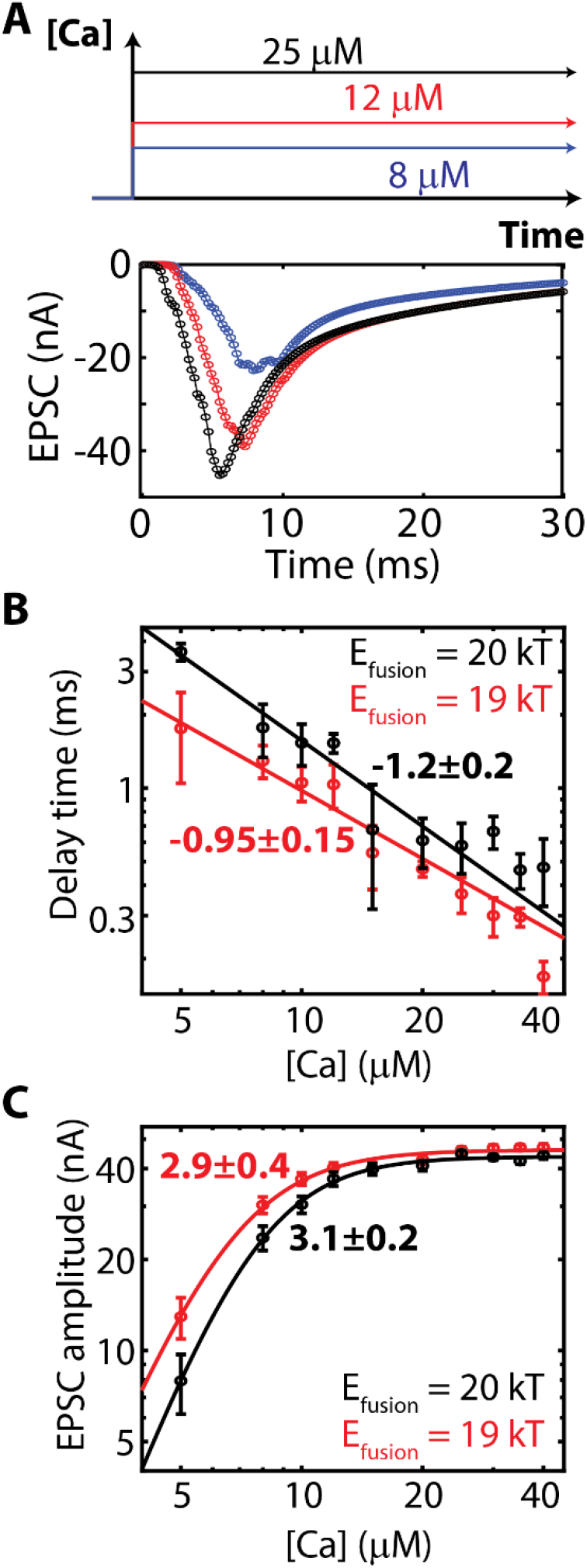
Increasing [Ca^2+^] increases EPSC amplitude and decreases synaptic delay. (A) Simulated and experimental Ca^2+^ uncaging-evoked EPSCs. (B) Synaptic delay times vs [Ca^2+^] in Ca^2+^-uncaging simulations with indicated energy barrier to fusion. Solid lines: best fit power laws, with indicated exponents. (C) EPSC amplitude vs [Ca^2+^] in Ca^2+^ uncaging simulations, with best-fit Hill functions and indicated Hill coefficients.

With increasing [Ca^2+^] unclamping and SNAREpin ring assembly became faster, and delay times decreased from 5.7±1.3 ms at 5 μM to 0.5±0.1 ms at 25 μM with a power law decay ~[Ca]^−1.2^, Fig. 5B, in quantitative agreement with the ~[Ca]^−1^ dependence observed in experiment of ref. (Lou et al., 2005).

In studies of the calyx of Held the delay time was only weakly affected when the fusion activation barrier was lowered by application of 1 μM phorbol ester phorbol-12,13-dibutyrate (PDBu) (Lou et al., 2005; Schotten et al., 2015). This is surprising, since significantly faster fusion would be expected to significantly decrease delays. Thus, we mimicked the effect of PDBu by lowering the fusion barrier to *E*_fusion_ = 19 *kT*. Fusion was ~2.6 times faster in simulations with SNAREs only, Fig. S5, but the synaptic delay time decreased by only 30-50%, Fig. 5B, consistent with these experiments.

The insensitivity of the delay time to the fusion rate is because the delay time is set by the very earliest release events. Since fusion times are exponentially distributed, Fig. S5, these earliest events occur immediately after unclamping, i.e. the fusion time is almost zero. Thus, the delay time is set primarily by the Ca^2+^-mediated unclamping time and is highly sensitive to presynaptic [Ca^2+^] but weakly dependent on the mean SNARE-mediated fusion rate.

### Dependence of EPSC amplitude on [Ca^2+^] does not reflect binding of 4 Ca^2+^ ions

It is hypothesized that four Ca^2+^ ions are required to trigger release of a vesicle (Augustine and Charlton, 1986; Rahamimoff and Dodge, 1969), based on the widely observed apparent cooperativity of ~3-5 for release, reviewed in (Neher and Sakaba, 2008).

Ca^2+^ uncaging simulations reproduced the experimental dependence of EPSC amplitude on [Ca^2+^], Fig. 5C. Fitting the amplitude *A* of the simulated EPSCs to a Hill function, 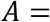 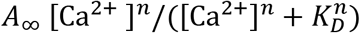, where *A*_∞_, *n*, and *K*_*D*_ are fitting parameters, yielded a Hill coefficient *n* = 3.1 ± 0.2 (the cooperativity) and apparent dissociation constant *KD* = 8.2 ± 0.3 μM, Fig. 5C.

Clearly, the apparent cooperativity of 3 in simulations is not directly related to the number of Ca^2+^ ions that trigger fusion, since ~30 Ca^2+^ ions were needed to unclamp each vesicle. Thus, the apparent cooperativity does not represent the number of Ca^2+^ ions needed for release, but is an emergent property of a complex, many-component machinery.

Lowering the energy threshold for fusion from 20 *kT* to 19 *kT* decreased the apparent dissociation constant to *K*_*D*_ = 7.4 ± 0.4 μM and increased the asymptotic EPSC amplitude, with almost no effect on the apparent cooperativity, Fig. 5C, reproducing the qualitative effect observed experimentally in ref. (Lou et al., 2005).

The plateau at high [Ca^2+^] in simulations, Fig. 5C, is a common feature of experimental dose-response curves (Heidelberger et al., 1994; Wang et al., 2008). The origin of the plateau was revealed by uncaging simulations starting from a fully unclamped state, when EPSC amplitudes were close to the asymptotic amplitudes in standard Ca^2+^ uncaging simulations, including a similar response to altered fusion barriers, Fig. S6. We conclude that the plateau is due to the Ca^2+^-dependent steps of NT release becoming so fast at high [Ca^2+^] that SNARE-mediated fusion becomes rate limiting.

### Action potentials evoke transient unclamping and low release probabilities

Next, we simulated physiological NT release at the calyx of Held and cerebellar granule cells, when [Ca^2+^] is elevated for just a brief ~1 ms window following an action potential (AP). We addressed an unexplained observation, that release probabilities are increased in autaptic hippocampal cultures by application of PDBu (Basu et al., 2007), thought to decrease the fusion barrier.

The [Ca^2+^] time dependence was taken from inferred Ca^2+^ transients at the calyx of Held based on a kinetic model of NT release (Wang et al., 2008), or from pre-synaptic currents *I*_Ca_ measured in cerebellar granule cells (Kawaguchi and Sakaba, 2017). EPSCs were obtained using experimental mEPSCs at the respective synapses (Malagon et al., 2016; Schneggenburger and Neher, 2000) (see Model section). For calyx of Held simulations, rate constants for Ca^2+^ binding/unbinding to C2B were fixed by the measured delay time and release probability (Schneggenburger and Neher, 2000), while for cerebellar granule cells the simulated peak [Ca^2+^] value was fixed by the release probability (Baur et al., 2015; Kawaguchi and Sakaba, 2017), Figs. S1 and S7 and Table S1.

During the course of the AP-evoked [Ca^2+^] transient, vesicles at both synapses were transiently unclamped (i.e. seven or more SNAREpins dissociated from the Syt ring). The unclamped fraction peaked at ~30% after ~0.25 ms at the calyx of Held (*n* = 1000), and at ~15% after ~1 ms in the cerebellar granule cell (*n* = 500), close to the [Ca^2+^] transient peak, Fig. 6A, C. Unreleased vesicles became re-clamped after [Ca^2+^] decreased to basal levels, so that ultimately only 10±1% or 18 ± 2% of vesicles released at the calyx of Held or cerebellar granule cells, respectively, Fig. 6B, D.

**Figure 6:**
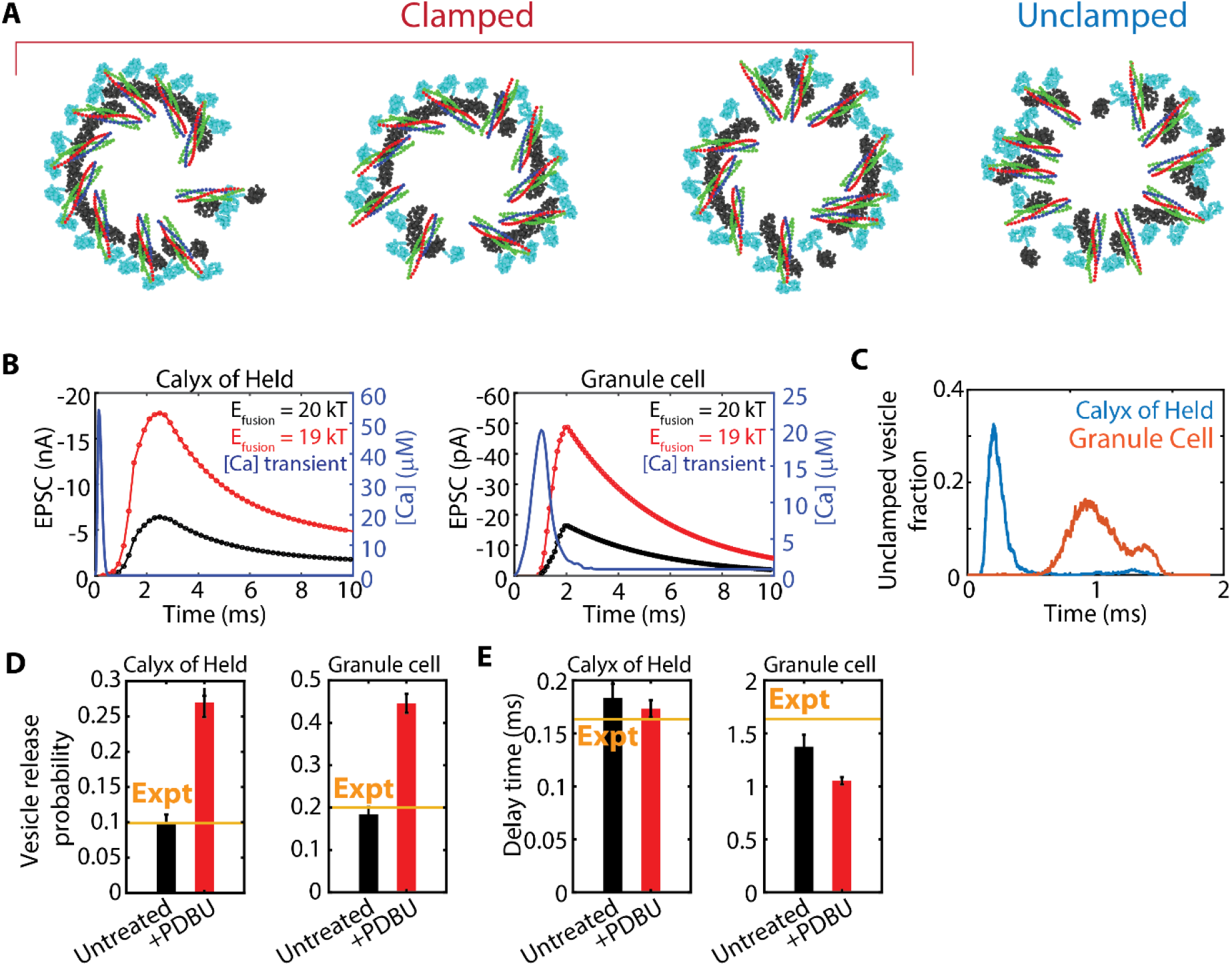
Action potentials evoke low release probabilities due to transient unclamping. (A) Snapshots from AP simulations at the calyx of Held, at the instant of peak [Ca^2+^] (~0.25 ms). (B) [Ca^2+^] transients (blue) and simulated EPSCs (red, black) at the calyx of Held (left, *n* = 1000 runs) and in cerebellar granule cells (center, *n* = 500 runs). (C) Fraction of vesicles that are unclamped vs time in the simulations of (B). (D) Vesicle release probabilities for the simulations of (B). Black (red) columns correspond to a fusion barrier of 20 *kT* (19 *kT*). Experimental values from (Kawaguchi and Sakaba, 2017; Schneggenburger and Neher, 2000). (E) Synaptic delay times for the simulations of (B). Experimental values from (Kawaguchi and Sakaba, 2017; Schneggenburger and Neher, 2000).

At both synapses, mimicking the effects of PDBu by lowering the fusion barrier to *E*_fusion_ = 19 *kT* from 20 *kT* decreased the synaptic delay time by only 20-30%, Fig. 6E, similarly to Ca^2+^ uncaging simulations. By contrast the release probability increased ~3-fold to ~0.26 and ~0.44 at the calyx of Held and cerebellar granule cell, respectively, consistent with increased release probabilities measured at hippocampal neurons treated with PDBu (Basu et al., 2007).

The simulations showed that the increased release probability originated in the transient nature of unclamping: with a higher fusion rate, vesicles were more likely to fuse during the time window when they were unclamped.

## Discussion

### Ca^2+^-sensing and membrane-fusing components of the release machinery are overlapping

The neurotransmitter release machinery includes Synaptotagmin, SNARE proteins, Munc18, Munc13 and Complexin (Brunger et al., 2018a; Sudhof, 2013). These and other components coordinate to sense calcium, execute unclamping and fuse the vesicle and plasma membranes. Synaptic delays, vesicle release rates and release probabilities are often interpreted in terms of Ca^2+^-sensing, unclamping and regulation of local Ca^2+^ concentration by calcium channel positioning and buffers (Dittman and Ryan, 2019; Eggermann et al., 2011; Neher and Sakaba, 2008) with SNARE-mediated fusion viewed as a decoupled final step following unclamping.

The present work suggests calcium-mediated unclamping and membrane fusion are overlapping functions of the machinery, and molecular components often associated with one aspect in fact service both, Fig. 7. Unclamping is driven by self-assembly of SNAREs into a ring, Fig. 4, and the shedding of Syt C2B-bound SNAREpins following membrane insertion of C2B Ca^2+^-binding loops. The shedding is due to structural constraints imposed by the SNARE-C2B complex (Voleti et al., 2020), Fig. 3. While the vesicle release probability is well known to be regulated by Ca^2+^-mediated unclamping (Dittman and Ryan, 2019; Eggermann et al., 2011; Vyleta and Jonas, 2014), we suggest the fusion component of the machinery plays an equally important role. By controlling the instantaneous fusion rate, it sets the probability that release occurs during the narrow unclamping time window, Figs. 6, 7.

**Figure 7:**
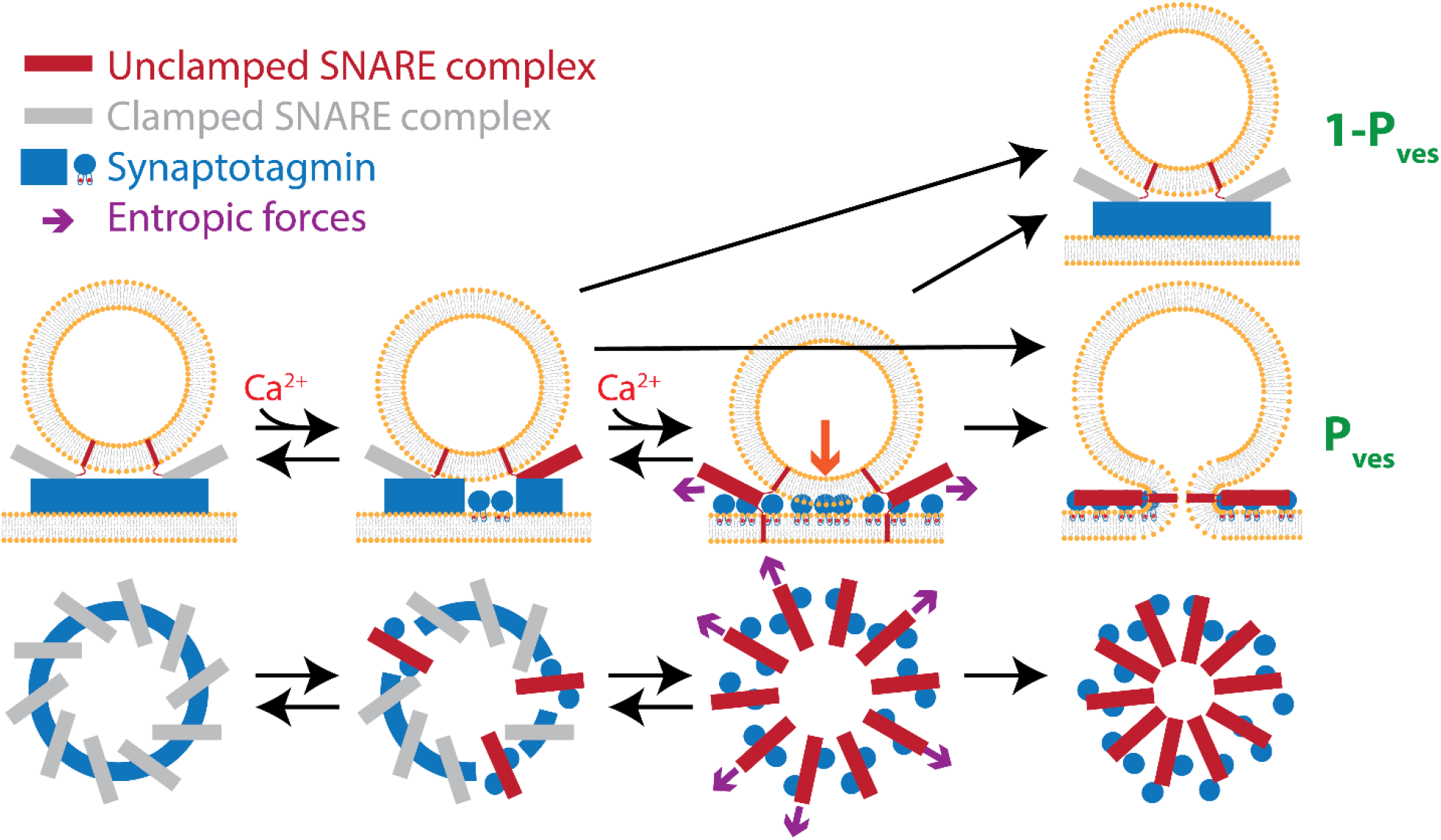
Model of coupled unclamping and fusion during action potential-evoked neurotransmitter release. Elevated [Ca^2+^] due to an action potential progressively unclamps SNARE complexes by dissociation from Synaptotagmin, driven by Ca^2+^-dependent insertion of [Ca^2+^]-binding loops of the C2B domain into the plasma membrane. The instantaneous fusion rate increases with the number of unclamped SNARE complexes, as more complexes exert higher entropic force and hence pull the membranes together with greater force (red arrow). Since unclamping is reversible and the elevated [Ca^2+^] is transient, the fate of a given vesicle is stochastic and can be fused or unfused. A fraction *P*_ves_ fuse, the release probability. Fusion can occur from a partially or completely unclamped state.

Other components of the release machinery may also overlap functionally with the Ca^2+^-sensing or membrane-fusing components. Several studies identified a “super-primed” vesicle state with elevated release probability associated with Munc13-Ca^2+^ interactions (Basu et al., 2007; Li et al., 2019a), and yeast vacuole fusion was proposed to be promoted by binding of SM-proteins to the SNARE complex N-terminus (D’Agostino et al., 2018).

### Membrane fusion is driven by entropic forces, not SNARE zippering energy

A common view is that fusion is driven by the SNARE complex zippering energy. The number of SNARE complexes required for fusion is then the fusion energy barrier divided by some fraction of the ~60 *kT* zippering energy (Gao et al., 2012; Ma et al., 2015). However, several observations challenge this notion. (i) No molecular machinery was suggested, to our knowledge, that could achieve this energy transduction, other than the original proposal that LD bending moments press the membranes together (Jahn and Scheller, 2006; Knecht and Grubmuller, 2003). However the SNARE LDs are likely unstructured and flexible (Kim et al., 2002; Lakomek et al., 2019), and insertion of helix-breaking proline residues progressively impairs but does not abolish fusion (Deak et al., 2006; McNew et al., 1999), although a recent study contradicts this claim (Hu et al., 2020). (ii) Fusion is often proposed triggered by calcium-induced release of SNARE complexes from a partially zippered state (Chicka et al., 2008; Gao et al., 2012; Giraudo et al., 2006). However, unrestrained zippering would be expected to require ~1 μs timescales (Gao et al., 2012; Kubelka et al., 2004), far less than the ~1 ms timescales of fusion (see below).

In our simulations, zippering on < 10 μs timescales was unrelated to fusion, other than to establish the bulky SNARE complexes. Fusion required a microscopically long delay, as fully zippered SNARE complexes pressed the membranes together in a waiting process with a constant fusion probability per unit time, Fig. S4. The pressing forces originated in entropic SNARE-SNARE and SNARE-membrane interactions that expanded the SNARE ring and pulled the membranes together, Fig. 5 & 7. We find fusion requires no minimum number of SNAREs, but rather is faster with more SNAREs since the entropic forces are then enhanced (McDargh et al., 2018; Mostafavi et al., 2017).

### Syt clamps release by spacing membranes and sequestering SNARE complexes

Release is evidently clamped before arrival of Ca^2+^, given the tiny spontaneous release rates per vesicle ~10^−6^ ms^−1^ (Lou et al., 2005) compared to evoked release rates, ~1 ms^−1^ (Wang et al., 2008). Syt knockout increases spontaneous release rates in mice and *Drosophila* (Geppert et al., 1994; Lee et al., 2013; Littleton et al., 1993), suggesting Syt is the clamp, possibly in concert with Complexin (Giraudo et al., 2006).

How Syt clamps release and then couples release to Ca^2+^ influx remains controversial. We assumed Syt oligomerizes into a ring at the fusion site, Fig. 1, consistent with impaired clamping in oligomerization-deficient Syt mutants (Bello et al., 2018; Tagliatti et al., 2020) and the abolition of a symmetric organization under docked vesicles in PC12 cells expressing the same mutant (Li et al., 2018). In simulations Syt rings clamped fusion by spacing the membranes, and by restraining the SNARE complexes bound to the ring, preventing their self-assembly into a ring that entropically presses the membranes together, Fig. 2. Consistent with the Syt-SNARE primary interaction contributing to clamping, spontaneous release frequencies are increased when the binding interface is disrupted but unaltered by mutations that strengthen it (Voleti et al., 2020; Zhou et al., 2015). In simulations, Ca^2+^ entry triggered ring disassembly and release of SNAREs by Ca^2+^-dependent membrane insertion of the C2B Ca^2+^-binding loops that sequestered the C2B domains, Fig. 3. The membranes could then meet, and the unfettered SNAREs could exert entropic forces and catalyze fusion.

Other models have been proposed. Many identify clamping with inhibition of SNARE complex assembly, directly removed by Ca^2+^ binding to Syt (Chicka et al., 2008; Grushin et al., 2019) or by displacing complexin (Zhou et al., 2017). Other models suggest Ca^2+^-dependent insertion of Syt Ca^2+^-binding loops into the PM induces curvature that facilitates fusion (Martens et al., 2007), or that Ca^2+^ binding provokes a conformational change in Syt with *trans* insertion of Ca^2+^-binding loops on C2A and C2B into the vesicle and plasma membranes, respectively (Nyenhuis et al., 2019; van den Bogaart et al., 2011).

### Unclamping progresses continuously from fully clamped to fully unclamped states

Lou et al. measured Ca^2+^ uncaging-evoked vesicle release rates with much lower cooperativities at low [Ca^2+^], and found that PDBu increased vesicle release rates and decreased delay times more weakly at higher [Ca^2+^], Figs. 5B, C (Lou et al., 2005). These findings are inconsistent with a sequential picture in which release requires a certain number of [Ca^2+^] ions to bind a single calcium sensor followed by a final fusion step. They devised a model of Ca^2+^-evoked NT release in which the instantaneous fusion rate depends on the number of Ca^2+^ ions bound to the Ca^2+^ sensor, assuming a maximum of 5 ions can bind. The model successfully reproduced these [Ca^2+^]-dependencies of the apparent Ca^2+^ cooperativity of release and sensitivity to PDBu, but did not attempt a molecular explanation.

Our simulations build on this phenomenological model by providing a molecularly explicit account of these effects, including the scaling of synaptic delay time with [Ca^2+^], the cooperativity of release at high [Ca^2+^], the increased release rates and reduced delay times on treatment with phorbol esters, and the weaking of this effect at higher [Ca^2+^], Figs. 5B, C. Realistic Ca^2+^ cooperativity values are predicted, without any assumptions on the number of Ca^2+^ ions needed to trigger release.

When [Ca^2+^] was elevated in simulations, the number of unclamped SNARE complexes increased gradually and reversibly, Figs. 3B & 7. As more SNARE complexes became activated by dissociation from the Syt ring, the instantaneous fusion rate increased. During an action potential, the degree to which a typical vesicle was unclamped peaked during the Ca^2+^ transient, and then declined after [Ca^2+^] returned to basal levels, Fig. 6C & 7.

### The importance of protein flexibility and fluctuations

Protein structures from cryo-EM or X-ray crystallography might suggest a static picture, but in reality structures fluctuate about the minimum energy state. Structural flexibility is important for drug discovery as drug targets may assume a range of conformational forms with similar energies (Teague, 2003), and for cellular processes such as membrane remodeling (Mahmood et al., 2019) and actin filament severing (McCullough et al., 2008).

We found that the flexibility of Syt rings is important to the pre-calcium clamped state. Simulated SNARE complexes bound to Syt rings were fully zippered, Fig. 3, in contrast to a proposal that zippering beyond layer +5 is inhibited by the binding of the SNARE complex to the C2B domain of Syt, based on cryo-EM measurements of the membrane-bound Syt-SNARE complex (Grushin et al., 2019). This inhibitory effect was reproduced only when simulated Syt rings were assumed perfectly rigid. Using the experimentally determined flexibility (Zhu et al., 2021), the rings could easily twist and accommodate complete zippering, Fig. 3. We conclude that constraints provided by Syt-SNARE binding alone are insufficient to impose a partially zippered state. However, we stress that our minimal model excluded other components such as Complexin that might inhibit C-terminal zippering by other means (Giraudo et al., 2006; Huntwork and Littleton, 2007). Indeed, evidence indicates that before Ca^2+^ influx the SNARE complexes are in a partially zippered state (Hua and Charlton, 1999; Xu et al., 1999), a feature absent from our model that presumably would require inclusion of additional components.

### Apparent cooperativity of release does not reflect binding of four Ca^2+^ ions to Syt

It has long been known that the post-synaptic current amplitude scales as the ~3-4^th^ power of [Ca^2+^] (the cooperativity), widely interpreted as indicating that ~4 Ca^2+^ ions are needed to trigger release (Augustine and Charlton, 1986; Katz and Miledi, 1970; Rahamimoff and Dodge, 1969; Schneggenburger and Neher, 2000). It was suggested this might correspond to the five Ca^2+^ ions found to bind the two C2 domains of Syt (Brose et al., 1992; Ubach et al., 1998).

With no fitting parameters, our simulations reproduced the experimental apparent cooperativity of release (Neher and Sakaba, 2008) but suggest the apparent ~4^th^ power is an emergent property of the complex machinery unrelated to the number of binding ions, as 12-40 ions were needed to unclamp each vesicle. There is likely no true power law scaling.

### Synaptic delay time is dominated by Ca^2+^-mediated unclamping

Long ago the synaptic delay between the AP peak to the onset of the EPSC at the frog neuromuscular junction was identified as the neurotransmitter release time, with small contributions from other processes (Katz and Miledi, 1965).

In simulations we found the synaptic delay is dominated by a subset of the release process, the time for Ca^2+^-mediated unclamping, Figs. 5 & 6. Since SNARE-mediated fusion is a quasi-steady state process, with fully assembled SNAREs imposing a constant fusion probability per unit time, the most probable fusion time is nearly zero, Fig. S5 (McDargh et al., 2018). Thus the synaptic delay measures the unclamping time with little contribution from fusion, being set by these earliest fusing vesicles. This explains the experimental observation that the delay time is relatively insensitive to the activation barrier for fusion (Lou et al., 2005) and the number of SNARE complexes at the fusion site (Acuna et al., 2014).

### SNARE-mediated fusion limits the neurotransmitter release rate at high [Ca^2+^]

Simulations reproduced the experimentally observed plateau in EPSC amplitude or vesicle release rate vs [Ca^2+^] at high concentrations (Bollmann et al., 2000; Schneggenburger and Neher, 2000; Wang et al., 2008), Figs. 5C & S8.

The plateau was due to SNARE-mediated fusion becoming rate-limiting at high [Ca^2+^]. This explains the observed decreases in asymptotic release rates at high [Ca^2+^] in hippocampal neurons expressing lower Syntaxin levels (Arancillo et al., 2013) or at presynaptic terminals at the calyx of Held treated with botulism and tetanus neurotoxins (Sakaba et al., 2005). In both cases we expect the fusion rate is lowered as the number of effective SNAREs at the fusion site is presumably lowered. However, we note that in other experiments a constitutively open Syntaxin mutant that increased the number of SNARE complexes at the fusion site did not elevate the asymptotic fusion rate (Acuna et al., 2014).

### Reading out unclamping and fusion times from experimental data

Overall, our model results suggest synaptic delay time at a given Ca^2+^ concentration is a readout of the unclamping time, while the asymptotic release rate per vesicle reads out the SNARE-mediated fusion rate. (1) Reported asymptotic release rates thus imply a fusion rate of ~0.5 − 1 ms^−1^ (Acuna et al., 2014; Sakaba et al., 2005; Schneggenburger and Neher, 2000; Wang et al., 2008), much slower than characteristic SNARE zippering rates, ~1 μs^−1^, supporting the view that zippering does not drive membrane fusion. (2) Measurements of synaptic delay times at the calyx of Held imply unclamping times ~0.5-1 ms for a presynaptic Ca^2+^ concentration of 25 μM, roughly the peak value at the release site during AP-evoked release (Bollmann et al., 2000; Schneggenburger and Neher, 2000).

Note that both the unclamping and fusion time exceed the typical width of local [Ca^2+^] transients during an AP at the calyx of Held (~0.25-0.5 ms (Bollmann et al., 2000; Borst and Sakmann, 1998; Wang et al., 2008)). This suggests most vesicles will fail to unclamp during the AP, consistent with the ~10% release probability (Schneggenburger and Neher, 2000).

### The SNARE-mediated fusion rate determines release probability

One might imagine vesicle release occurs if and only if Ca^2+^-mediated unclamping occurs, so the release probability is unrelated to fusion. In the framework of a Syt ring-based model of the NT machinery, Fig. 1, simulations showed this is far from true, since unclamping is reversible: during an action potential, many vesicles were unclamped for a short window of time during the [Ca^2+^] transient, Fig. 6C & 7. As Ca^2+^ ions dissociated from Syt monomers, vesicles became reclamped as Syt monomers reoligomerized and reformed bonds with SNARE complexes, re-sequestering them and inhibiting the entropic forces that drive fusion. Thus the net vesicle release probability depends on the fusion probability during the finite unclamping window, so that higher fusion rates drive higher release probabilities, consistent with the observed increased of release probabilities following PDBu treatment (Basu et al., 2007; Lou et al., 2005; Schotten et al., 2015).

This prediction is also supported by studies with decreased (Arancillo et al., 2013) or increased (Acuna et al., 2014) numbers of SNARE complexes mediating release, when release probabilities were decreased or increased, respectively. This accords with our predictions, as more SNAREs give higher fusion rates (McDargh et al., 2018; Mostafavi et al., 2017). In a similar vein, we speculate that lowered vesicle release probabilities induced by Complexin knockout (Xue et al., 2010) or altered SNARE complex charge (Ruiter et al., 2019) may originate in decreased fusion rates.

## References

Acuna, C., Guo, Q., Burré, J., Sharma, M., Sun, J., and Südhof, T.C. (2014). Microsecond Dissection of Neurotransmitter Release: SNARE-Complex Assembly Dictates Speed and Ca2+ Sensitivity. Neuron 82, 1088–1100.

Arancillo, M., Min, S.W., Gerber, S., Munster-Wandowski, A., Wu, Y.J., Herman, M., Trimbuch, T., Rah, J.C., Ahnert-Hilger, G., Riedel, D., et al. (2013). Titration of Syntaxin1 in mammalian synapses reveals multiple roles in vesicle docking, priming, and release probability. J Neurosci 33, 16698–16714.

Augustine, G.J., and Charlton, M.P. (1986). Calcium dependence of presynaptic calcium current and post-synaptic response at the squid giant synapse. J Physiol 381, 619–640.

Basu, J., Betz, A., Brose, N., and Rosenmund, C. (2007). Munc13-1 C1 domain activation lowers the energy barrier for synaptic vesicle fusion. J Neurosci 27, 1200–1210.

Baur, D., Bornschein, G., Althof, D., Watanabe, M., Kulik, A., Eilers, J., and Schmidt, H. (2015). Developmental tightening of cerebellar cortical synaptic influx-release coupling. J Neurosci 35, 1858–1871.

Bello, O.D., Jouannot, O., Chaudhuri, A., Stroeva, E., Coleman, J., Volynski, K.E., Rothman, J.E., and Krishnakumar, S.S. (2018). Synaptotagmin oligomerization is essential for calcium control of regulated exocytosis. Proc Natl Acad Sci U S A 115, E7624–E7631.

Bohme, M.A., Grasskamp, A.T., and Walter, A.M. (2018). Regulation of synaptic release-site Ca(2+) channel coupling as a mechanism to control release probability and short-term plasticity. FEBS Lett 592, 3516–3531.

Bollmann, J.H., Sakmann, B., and Borst, J.G. (2000). Calcium sensitivity of glutamate release in a calyx-type terminal. Science 289, 953–957.

Borst, J.G., and Sakmann, B. (1998). Calcium current during a single action potential in a large presynaptic terminal of the rat brainstem. J Physiol 506 (Pt 1), 143–157.

Branco, T., and Staras, K. (2009). The probability of neurotransmitter release: variability and feedback control at single synapses. Nat Rev Neurosci 10, 373–383.

Broadie, K., Bellen, H.J., DiAntonio, A., Littleton, J.T., and Schwarz, T.L. (1994). Absence of synaptotagmin disrupts excitation-secretion coupling during synaptic transmission. Proc Natl Acad Sci U S A 91, 10727–10731.

Brose, N., Petrenko, A.G., Sudhof, T.C., and Jahn, R. (1992). Synaptotagmin: a calcium sensor on the synaptic vesicle surface. Science 256, 1021–1025.

Brunger, A.T., Choi, U.B., Lai, Y., Leitz, J., and Zhou, Q. (2018a). Molecular Mechanisms of Fast Neurotransmitter Release. Annu Rev Biophys 47, 469–497.

Brunger, A.T., Leitz, J., Zhou, Q., Choi, U.B., and Lai, Y. (2018b). Ca(2+)-Triggered Synaptic Vesicle Fusion Initiated by Release of Inhibition. Trends Cell Biol 28, 631–645.

Burgalossi, A., Jung, S., Meyer, G., Jockusch, W.J., Jahn, O., Taschenberger, H., O’Connor, V.M., Nishiki, T., Takahashi, M., Brose, N., et al. (2010). SNARE protein recycling by alphaSNAP and betaSNAP supports synaptic vesicle priming. Neuron 68, 473–487.

Chicka, M.C., Hui, E., Liu, H., and Chapman, E.R. (2008). Synaptotagmin arrests the SNARE complex before triggering fast, efficient membrane fusion in response to Ca2+. Nat Struct Mol Biol 15, 827–835.

D’Agostino, M., Risselada, H.J., Endter, L.J., Comte-Miserez, V., and Mayer, A. (2018). SNARE-mediated membrane fusion arrests at pore expansion to regulate the volume of an organelle. EMBO J 37.

Deak, F., Shin, O.H., Kavalali, E.T., and Sudhof, T.C. (2006). Structural determinants of synaptobrevin 2 function in synaptic vesicle fusion. J Neurosci 26, 6668–6676.

Dittman, J.S., and Ryan, T.A. (2019). The control of release probability at nerve terminals. Nat Rev Neurosci 20, 177–186.

Eggermann, E., Bucurenciu, I., Goswami, S.P., and Jonas, P. (2011). Nanodomain coupling between Ca(2)(+) channels and sensors of exocytosis at fast mammalian synapses. Nat Rev Neurosci 13, 7–21.

Ehlers, M.D., Mammen, A.L., Lau, L.F., and Huganir, R.L. (1996). Synaptic targeting of glutamate receptors. Curr Opin Cell Biol 8, 484–489.

Ermolyuk, Y.S., Alder, F.G., Surges, R., Pavlov, I.Y., Timofeeva, Y., Kullmann, D.M., and Volynski, K.E. (2013). Differential triggering of spontaneous glutamate release by P/Q-, N- and R-type Ca2+ channels. Nat Neurosci 16, 1754–1763.

Fernandez-Chacon, R., Konigstorfer, A., Gerber, S.H., Garcia, J., Matos, M.F., Stevens, C.F., Brose, N., Rizo, J., Rosenmund, C., and Sudhof, T.C. (2001). Synaptotagmin I functions as a calcium regulator of release probability. Nature 410, 41–49.

Fernandez, I., Arac, D., Ubach, J., Gerber, S.H., Shin, O., Gao, Y., Anderson, R.G., Sudhof, T.C., and Rizo, J. (2001). Three-dimensional structure of the synaptotagmin 1 C2B-domain: synaptotagmin 1 as a phospholipid binding machine. Neuron 32, 1057–1069.

Fioravante, D., and Regehr, W.G. (2011). Short-term forms of presynaptic plasticity. Curr Opin Neurobiol 21, 269–274.

Gao, Y., Zorman, S., Gundersen, G., Xi, Z., Ma, L., Sirinakis, G., Rothman, J.E., and Zhang, Y. (2012). Single reconstituted neuronal SNARE complexes zipper in three distinct stages. Science 337, 1340–1343.

Geppert, M., Goda, Y., Hammer, R.E., Li, C., Rosahl, T.W., Stevens, C.F., and Sudhof, T.C. (1994). Synaptotagmin I: a major Ca2+ sensor for transmitter release at a central synapse. Cell 79, 717–727.

Giraudo, C.G., Eng, W.S., Melia, T.J., and Rothman, J.E. (2006). A clamping mechanism involved in SNARE-dependent exocytosis. Science 313, 676–680.

Grushin, K., Wang, J., Coleman, J., Rothman, J.E., Sindelar, C.V., and Krishnakumar, S.S. (2019). Structural basis for the clamping and Ca(2+) activation of SNARE-mediated fusion by synaptotagmin. Nat Commun 10, 2413.

Heidelberger, R., Heinemann, C., Neher, E., and Matthews, G. (1994). Calcium dependence of the rate of exocytosis in a synaptic terminal. Nature 371, 513–515.

Hu, Y., Zhu, L., and Ma, C. (2020). Structural Roles for the Juxtamembrane Linker Region and Transmembrane Region of Synaptobrevin 2 in Membrane Fusion. Front Cell Dev Biol 8, 609708.

Hua, S.Y., and Charlton, M.P. (1999). Activity-dependent changes in partial VAMP complexes during neurotransmitter release. Nat Neurosci 2, 1078–1083.

Huntwork, S., and Littleton, J.T. (2007). A complexin fusion clamp regulates spontaneous neurotransmitter release and synaptic growth. Nat Neurosci 10, 1235–1237.

Jackson, M.B., and Redman, S.J. (2003). Calcium dynamics, buffering, and buffer saturation in the boutons of dentate granule-cell axons in the hilus. J Neurosci 23, 1612–1621.

Jahn, R., and Scheller, R.H. (2006). SNAREs - engines for membrane fusion. Nature Reviews Molecular Cell Biology 7, 631–643.

Katz, B., and Miledi, R. (1965). The Measurement of Synaptic Delay, and the Time Course of Acetylcholine Release at the Neuromuscular Junction. Proc R Soc Lond B Biol Sci 161, 483–495.

Kawaguchi, S.Y., and Sakaba, T. (2017). Fast Ca(2+) Buffer-Dependent Reliable but Plastic Transmission at Small CNS Synapses Revealed by Direct Bouton Recording. Cell Rep 21, 3338–3345.

Kim, C.S., Kweon, D.H., and Shin, Y.K. (2002). Membrane topologies of neuronal SNARE folding intermediates. Biochemistry 41, 10928–10933.

Knecht, V., and Grubmuller, H. (2003). Mechanical coupling via the membrane fusion SNARE protein syntaxin 1A: a molecular dynamics study. Biophys J 84, 1527–1547.

Korber, C., and Kuner, T. (2016). Molecular Machines Regulating the Release Probability of Synaptic Vesicles at the Active Zone. Front Synaptic Neurosci 8, 5.

Kubelka, J., Hofrichter, J., and Eaton, W.A. (2004). The protein folding ‘speed limit’. Curr Opin Struct Biol 14, 76–88.

Lai, Y., Choi, U.B., Leitz, J., Rhee, H.J., Lee, C., Altas, B., Zhao, M., Pfuetzner, R.A., Wang, A.L., Brose, N., et al. (2017). Molecular Mechanisms of Synaptic Vesicle Priming by Munc13 and Munc18. Neuron 95, 591–607 e510.

Lakomek, N.A., Yavuz, H., Jahn, R., and Perez-Lara, A. (2019). Structural dynamics and transient lipid binding of synaptobrevin-2 tune SNARE assembly and membrane fusion. Proc Natl Acad Sci U S A 116, 8699–8708.

Lee, J., Guan, Z., Akbergenova, Y., and Littleton, J.T. (2013). Genetic analysis of synaptotagmin C2 domain specificity in regulating spontaneous and evoked neurotransmitter release. J Neurosci 33, 187–200.

Li, L., Liu, H., Hall, Q., Wang, W., Yu, Y., Kaplan, J.M., and Hu, Z. (2019a). A Hyperactive Form of unc-13 Enhances Ca(2+) Sensitivity and Synaptic Vesicle Release Probability in C. elegans. Cell Rep 28, 2979–2995 e2974.

Li, X., Radhakrishnan, A., Grushin, K., Kasula, R., Chaudhuri, A., Gomathinayagam, S., Krishnakumar, S.S., Liu, J., and Rothman, J.E. (2018). Symmetrical Organization of Proteins Under Docked Synaptic-Vesicles. FEBS Lett.

Li, X., Radhakrishnan, A., Grushin, K., Kasula, R., Chaudhuri, A., Gomathinayagam, S., Krishnakumar, S.S., Liu, J., and Rothman, J.E. (2019b). Symmetrical organization of proteins under docked synaptic vesicles. FEBS Lett 593, 144–153.

Littleton, J.T., Stern, M., Perin, M., and Bellen, H.J. (1994). Calcium dependence of neurotransmitter release and rate of spontaneous vesicle fusions are altered in Drosophila synaptotagmin mutants. Proc Natl Acad Sci U S A 91, 10888–10892.

Littleton, J.T., Stern, M., Schulze, K., Perin, M., and Bellen, H.J. (1993). Mutational analysis of Drosophila synaptotagmin demonstrates its essential role in Ca(2+)-activated neurotransmitter release. Cell 74, 1125–1134.

Lou, X., Scheuss, V., and Schneggenburger, R. (2005). Allosteric modulation of the presynaptic Ca2+ sensor for vesicle fusion. Nature 435, 497–501.

Ma, L., Cai, Y., Li, Y., Jiao, J., Wu, Z., O’Shaughnessy, B., De Camilli, P., Karatekin, E., and Zhang, Y. (2017). Single-molecule force spectroscopy of protein-membrane interactions. Elife 6.

Ma, L., Rebane, A.A., Yang, G., Xi, Z., Kang, Y., Gao, Y., and Zhang, Y. (2015). Munc18-1-regulated stage-wise SNARE assembly underlying synaptic exocytosis. Elife 4.

Mahmood, M.I., Noguchi, H., and Okazaki, K.I. (2019). Curvature induction and sensing of the F-BAR protein Pacsin1 on lipid membranes via molecular dynamics simulations. Sci Rep 9, 14557.

Malagon, G., Miki, T., Llano, I., Neher, E., and Marty, A. (2016). Counting Vesicular Release Events Reveals Binomial Release Statistics at Single Glutamatergic Synapses. J Neurosci 36, 4010–4025.

Martens, S., Kozlov, M.M., and McMahon, H.T. (2007). How synaptotagmin promotes membrane fusion. Science 316, 1205–1208.

McCullough, B.R., Blanchoin, L., Martiel, J.L., and De la Cruz, E.M. (2008). Cofilin increases the bending flexibility of actin filaments: implications for severing and cell mechanics. J Mol Biol 381, 550–558.

McDargh, Z.A., Polley, A., and O’Shaughnessy, B. (2018). SNARE-mediated membrane fusion is a two-stage process driven by entropic forces. FEBS Lett 592, 3504–3515.

McNew, J.A., Weber, T., Engelman, D.M., Sollner, T.H., and Rothman, J.E. (1999). The length of the flexible SNAREpin juxtamembrane region is a critical determinant of SNARE-dependent fusion. Molecular Cell 4, 415–421.

Mostafavi, H., Thiyagarajan, S., Stratton, B.S., Karatekin, E., Warner, J.M., Rothman, J.E., and O’Shaughnessy, B. (2017). Entropic forces drive self-organization and membrane fusion by SNARE proteins. Proc Natl Acad Sci U S A 114, 5455–5460.

Neher, E., and Sakaba, T. (2008). Multiple roles of calcium ions in the regulation of neurotransmitter release. Neuron 59, 861–872.

Nishiki, T., and Augustine, G.J. (2004a). Dual roles of the C2B domain of synaptotagmin I in synchronizing Ca2+-dependent neurotransmitter release. J Neurosci 24, 8542–8550.

Nishiki, T., and Augustine, G.J. (2004b). Synaptotagmin I synchronizes transmitter release in mouse hippocampal neurons. J Neurosci 24, 6127–6132.

Nyenhuis, S.B., Thapa, A., and Cafiso, D.S. (2019). Phosphatidylinositol 4,5 Bisphosphate Controls the cis and trans Interactions of Synaptotagmin 1. Biophys J 117, 247–257.

Perez-Lara, A., Thapa, A., Nyenhuis, S.B., Nyenhuis, D.A., Halder, P., Tietzel, M., Tittmann, K., Cafiso, D.S., and Jahn, R. (2016). PtdInsP2 and PtdSer cooperate to trap synaptotagmin-1 to the plasma membrane in the presence of calcium. Elife 5.

Radhakrishnan, A., Stein, A., Jahn, R., and Fasshauer, D. (2009). The Ca2+ affinity of synaptotagmin 1 is markedly increased by a specific interaction of its C2B domain with phosphatidylinositol 4,5-bisphosphate. J Biol Chem 284, 25749–25760.

Rahamimoff, R., and Dodge, F.A., Jr. (1969). Regulation of transmitter release at the neuromuscular synapse: the cooperative hypothesis. Electroencephalogr Clin Neurophysiol 27, 219.

Ramakrishnan, S., Bera, M., Coleman, J., Krishnakumar, S.S., Pincet, F., and Rothman, J.E. (2018). Synaptotagmin oligomers are necessary and can be sufficient to form a Ca(2+) -sensitive fusion clamp. FEBS Lett.

Ramakrishnan, S., Bera, M., Coleman, J., Rothman, J.E., and Krishnakumar, S.S. (2020). Synergistic roles of Synaptotagmin-1 and complexin in calcium-regulated neuronal exocytosis. Elife 9.

Rhee, J.S., Li, L.Y., Shin, O.H., Rah, J.C., Rizo, J., Sudhof, T.C., and Rosenmund, C. (2005). Augmenting neurotransmitter release by enhancing the apparent Ca2+ affinity of synaptotagmin 1. Proc Natl Acad Sci U S A 102, 18664–18669.

Rosenmund, C., Sigler, A., Augustin, I., Reim, K., Brose, N., and Rhee, J.S. (2002). Differential control of vesicle priming and short-term plasticity by Munc13 isoforms. Neuron 33, 411–424.

Rosenmund, C., and Stevens, C.F. (1996). Definition of the readily releasable pool of vesicles at hippocampal synapses. Neuron 16, 1197–1207.

Ruiter, M., Kadkova, A., Scheutzow, A., Malsam, J., Sollner, T.H., and Sorensen, J.B. (2019). An Electrostatic Energy Barrier for SNARE-Dependent Spontaneous and Evoked Synaptic Transmission. Cell Rep 26, 2340–2352 e2345.

Sabatini, B.L., and Regehr, W.G. (1996). Timing of neurotransmission at fast synapses in the mammalian brain. Nature 384, 170–172.

Sakaba, T. (2008). Two Ca(2+)-dependent steps controlling synaptic vesicle fusion and replenishment at the cerebellar basket cell terminal. Neuron 57, 406–419.

Sakaba, T., Stein, A., Jahn, R., and Neher, E. (2005). Distinct kinetic changes in neurotransmitter release after SNARE protein cleavage. Science 309, 491–494.

Schneggenburger, R., and Neher, E. (2000). Intracellular calcium dependence of transmitter release rates at a fast central synapse. Nature 406, 889–893.

Schotten, S., Meijer, M., Walter, A.M., Huson, V., Mamer, L., Kalogreades, L., ter Veer, M., Ruiter, M., Brose, N., Rosenmund, C., et al. (2015). Additive effects on the energy barrier for synaptic vesicle fusion cause supralinear effects on the vesicle fusion rate. Elife 4, e05531.

Sollner, T., Whiteheart, S.W., Brunner, M., Erdjument-Bromage, H., Geromanos, S., Tempst, P., and Rothman, J.E. (1993). SNAP receptors implicated in vesicle targeting and fusion. Nature 362, 318–324.

Stein, A., Weber, G., Wahl, M.C., and Jahn, R. (2009). Helical extension of the neuronal SNARE complex into the membrane. Nature 460, 525–U105.

Sudhof, T.C. (2013). Neurotransmitter release: the last millisecond in the life of a synaptic vesicle. Neuron 80, 675–690.

Sun, J., Pang, Z.P., Qin, D., Fahim, A.T., Adachi, R., and Sudhof, T.C. (2007). A dual-Ca2+-sensor model for neurotransmitter release in a central synapse. Nature 450, 676–682.

Tagliatti, E., Bello, O.D., Mendonca, P.R.F., Kotzadimitriou, D., Nicholson, E., Coleman, J., Timofeeva, Y., Rothman, J.E., Krishnakumar, S.S., and Volynski, K.E. (2020). Synaptotagmin 1 oligomers clamp and regulate different modes of neurotransmitter release. Proc Natl Acad Sci U S A 117, 3819–3827.

Takamori, S., Holt, M., Stenius, K., Lemke, E.A., Gronborg, M., Riedel, D., Urlaub, H., Schenck, S., Brugger, B., Ringler, P., et al. (2006). Molecular anatomy of a trafficking organelle. Cell 127, 831–846.

Teague, S.J. (2003). Implications of protein flexibility for drug discovery. Nat Rev Drug Discov 2, 527–541.

Ubach, J., Zhang, X., Shao, X., Sudhof, T.C., and Rizo, J. (1998). Ca2+ binding to synaptotagmin: how many Ca2+ ions bind to the tip of a C2-domain? EMBO J 17, 3921–3930.

van den Bogaart, G., Thutupalli, S., Risselada, J.H., Meyenberg, K., Holt, M., Riedel, D., Diederichsen, U., Herminghaus, S., Grubmuller, H., and Jahn, R. (2011). Synaptotagmin-1 may be a distance regulator acting upstream of SNARE nucleation. Nat Struct Mol Biol 18, 805–812.

Voleti, R., Jaczynska, K., and Rizo, J. (2020). Ca(2+)-dependent release of Synaptotagmin-1 from the SNARE complex on phosphatidylinositol 4,5-bisphosphate-containing membranes. Elife 9.

Vyleta, N.P., and Jonas, P. (2014). Loose coupling between Ca2+ channels and release sensors at a plastic hippocampal synapse. Science 343, 665–670.

Wang, J., Bello, O., Auclair, S.M., Wang, J., Coleman, J., Pincet, F., Krishnakumar, S.S., Sindelar, C.V., and Rothman, J.E. (2014). Calcium sensitive ring-like oligomers formed by synaptotagmin. Proc Natl Acad Sci U S A 111, 13966–13971.

Wang, J., Li, F., Bello, O.D., Sindelar, C.V., Pincet, F., Krishnakumar, S.S., and Rothman, J.E. (2017). Circular oligomerization is an intrinsic property of synaptotagmin. Elife 6.

Wang, L.Y., Neher, E., and Taschenberger, H. (2008). Synaptic vesicles in mature calyx of Held synapses sense higher nanodomain calcium concentrations during action potential-evoked glutamate release. J Neurosci 28, 14450–14458.

Weber, T., Zemelman, B.V., McNew, J.A., Westermann, B., Gmachl, M., Parlati, F., Sollner, T.H., and Rothman, J.E. (1998). SNAREpins: minimal machinery for membrane fusion. Cell 92, 759–772.

Wilhelm, B.G., Mandad, S., Truckenbrodt, S., Krohnert, K., Schafer, C., Rammner, B., Koo, S.J., Classen, G.A., Krauss, M., Haucke, V., et al. (2014). Composition of isolated synaptic boutons reveals the amounts of vesicle trafficking proteins. Science 344, 1023–1028.

Xu, T., Rammner, B., Margittai, M., Artalejo, A.R., Neher, E., and Jahn, R. (1999). Inhibition of SNARE complex assembly differentially affects kinetic components of exocytosis. Cell 99, 713–722.

Xue, M., Craig, T.K., Xu, J., Chao, H.T., Rizo, J., and Rosenmund, C. (2010). Binding of the complexin N terminus to the SNARE complex potentiates synaptic-vesicle fusogenicity. Nat Struct Mol Biol 17, 568–575.

Zanetti, M.N., Bello, O.D., Wang, J., Coleman, J., Cai, Y., Sindelar, C.V., Rothman, J.E., and Krishnakumar, S.S. (2016). Ring-like oligomers of Synaptotagmins and related C2 domain proteins. Elife 5.

Zhou, Q., Lai, Y., Bacaj, T., Zhao, M., Lyubimov, A.Y., Uervirojnangkoorn, M., Zeldin, O.B., Brewster, A.S., Sauter, N.K., Cohen, A.E., et al. (2015). Architecture of the synaptotagmin-SNARE machinery for neuronal exocytosis. Nature 525, 62–67.

Zhou, Q., Zhou, P., Wang, A.L., Wu, D., Zhao, M., Sudhof, T.C., and Brunger, A.T. (2017). The primed SNARE-complexin-synaptotagmin complex for neuronal exocytosis. Nature 548, 420–425.

Zhu, J., McDargh, Z.A., Li, F., Krishnakumar, S., Rothman, J.E., and O’Shaughnessy, B. (2021). Synaptotagmin rings as high sensitivity regulators of synaptic vesicle docking and fusion. bioRxiv, 2021.2003.2012.435193.

