## Supporting Information for "Coupling of Ca^2+^-triggered unclamping and membrane fusion during neurotransmitter release"

### <u>Supplementary Material</u>

#### Coarse-grained models

In order to achieve the long time-scales of NT release, our simulations used highly coarse-grained (CG) models of the SNARE complex, Synaptotagmin, and the vesicle and plasma membranes.

*SNARE complex:* The model of the SNARE complex used in our simulations was previously applied in similar simulations in refs (McDargh et al., 2018; Mostafavi et al., 2017). For completeness, we review the model's design here. As input, the model takes the all-atom crystal structure of the fully assembled SNARE complex found in reference (Stein et al., 2009) (PDB ID 3HD7). The SNARE complex is a four helix coiled-coil comprising 16 layers, numbered -7 to +8. Groups of 4 contiguous residues along the backbone of each helix are mapped to a single CG bead, corresponding roughly to one layer per CG bead.

In the model, the beads making up each layer are arranged into a square of side-length 1.33 nm, roughly the separation between the centers of the alpha helices making up the SNARE complex coiled-coil. The radius of each bead is 0.33 nm, reproducing the overall ~2 nm thickness of the SNARE complex. Adjacent layers are separated by 0.66 nm, reproducing the overall ~10 nm length of the SNARE complex. Each layer is rotated by 14° about the axis of the SNARE complex relative to the neighboring layers.

Uncomplexed layers of VAMP and Syntaxin are represented by worm-like chains, with contour length varying depending on the degree of assembly of the SNARE complex. The stretching energy of the worm-like chain is

$$E_{\text{linker}} = kT \frac{l_c}{l_p} \frac{1}{4\left(1 - \frac{r}{l_c}\right)} \left(3\left(\frac{r}{l_c}\right)^2 - 2\left(\frac{r}{l_c}\right)^3\right),$$

where $l_p$ =0.6 nm is the persistence length (a typical value for unstructured polypeptides (Gao et al., 2012; Gebhardt et al., 2010)), $r$ is the end-to-end distance of the unstructured segment, and $l_c$ is contour length, calculated from the total number of uncomplexed residues in the LD, assuming that each residue contributes 0.365 nm to the total contour length (Gao et al., 2012).

The magnitude force caused by stretching the linker is obtained by differentiating the energy with respect to $r$, and the force is directed from one endpoint of the linker towards the other. The Syntaxin linker connects from the last complexed bead of the Syntaxin helix to the closest point on the PM surface to that bead; similarly, the VAMP linker connects from the last complexed bead of the VAMP helix to the closest point on the surface of the vesicle to that bead.

*Synaptotagmin 1:* Our Synaptotagmin 1 model uses a similar method to that of the SNARE complex model. For reference, we take the crystal structure of the Syt C2AB fragment measured in reference (Zhou et al., 2015) (PDB ID 5CCG). Groups of contiguous residues along the protein backbone are mapped to a single CG bead: if the residues are part of an alpha helix or beta sheet, 4 residues are mapped to a bead, while if the residues are unstructured, 2 residues are mapped to a bead. This gives the model greater resolution in regions where the protein backbone is most highly curved. The 60-residue linker connecting the C2A domain to the trans-membrane domain is represented by a worm-like chain (discussed in **Supp. Mat.: Non-bonded interactions**). The trans-membrane domain is assumed to be located on the surface of the vesicle at the closest point to the first bead of the C2A domain (i.e. the bead at which the linker attaches).

Each CG bead is placed at the center of mass of its constituent residues. The radius of the bead is chosen to preserve the total van der Waals volume of its constituent beads.

*Membranes:* Membranes are represented as non-deformable continuous surfaces. The plasma membrane forms a plane at fixed position $z = 0$, while the vesicle is represented by a sphere of radius 20 nm. The PM composition is assumed to be 40% PE, 40% PC, 18% PS, and 2% PI (Hitzemann and Johnson, 1983), while the vesicle membrane is assumed to be 23% PE, 46% PC, and 12% PS (Takamori et al., 2006), Table S2.

**Molecular dynamics simulations**

*Translational motion:* The simulation procedure used here is identical to that used in our previous study, ref. (McDargh et al., 2018). We describe it here for completeness. The proteins and vesicles in our simulations obey the equations of Langevin dynamics in the over-damped regime, i.e. with no inertial term. The velocity of the center of mass of a protein or vesicle is determined at each time step as

$$\gamma \boldsymbol{V} = \boldsymbol{F} + \boldsymbol{\eta}$$

where $\gamma$ is the total drag coefficient of the protein or vesicle, $\boldsymbol{V}$ is the velocity of its center of mass, $\boldsymbol{F}$ is the vector sum of all the interaction forces on the protein's constituent beads (or on the vesicle as a whole), and $\boldsymbol{\eta}$ is the vector sum of all stochastic noise forces (discussed below) on the protein's constituent beads (or on the vesicle as a whole). The drag coefficient of each protein is the sum of the drag coefficient of its constituent beads, while the drag coefficient of each bead is determined by the Stokes formula $\gamma_{\mathrm{bead}} = 6\pi\nu R_{\mathrm{bead}}$, where $\nu$ is the viscosity of water (see Table S1) and $R_{\mathrm{bead}}$ is the radius of the bead.

*Rotational motion:* Similarly, the rotational velocity is determined at each time step as

$$\boldsymbol{\Gamma\Omega} = \boldsymbol{M} + \boldsymbol{\nu}$$

where $\boldsymbol{\Gamma}$ is the rotational drag matrix of the protein, $\boldsymbol{\Omega}$ is the angular velocity, $\boldsymbol{M}$ is the total moment of all interaction forces on the protein's constituent beads, and $\boldsymbol{\nu}$ is the total moment due to stochastic noise forces on the constituent beads. Using index notation, the rotational drag matrix of a protein is given by

$$\Gamma_{ij} = \sum_b (\delta_{ij} d_b^2 - d_{b_i} d_{b_j})\, \gamma_b,$$

where $\boldsymbol{d}_b$ is the vector pointing from the center of mass of the protein to the center of bead $b$, $\delta_{ij}$ is the Kronecker delta, and $\gamma_b$ is the drag coefficient of bead $b$, and the sum is taken over all beads of the protein.

The total moment of the interaction forces on the protein is the sum of the moments due to interactions of the protein's constituent beads,

$$\boldsymbol{M} = \sum_b \boldsymbol{f}_b \times \boldsymbol{d}_b, \ \ \boldsymbol{\nu} = \sum_b \boldsymbol{\eta}_b \times \boldsymbol{d}_b$$

where $\boldsymbol{f}_b$ is the net force due to interactions on bead $b$, and $\boldsymbol{\eta}_b$ is the noise force on bead $b$.

*Noise forces:* The stochastic noise forces are calculated on a bead-by-bead basis. At each time step, a random force is generated for each bead in the simulation such that the spatial components of the force are uniformly distributed over the interval $[-\sqrt{6\,\gamma_{\text{bead}}\,kT/\delta t}, +\sqrt{6\,\gamma_{\text{bead}}\,kT/\delta t}]$, where $k$ is the Boltzmann constant, $T$ is room temperature, and $\delta t$ is the time step of the simulation. Crucially, this guarantees that the force has zero mean, and variance $\langle {\eta_{b}}_{i}^{2} \rangle = 2\,\gamma_{\text{bead}} kT/\delta t$, thus setting the temperature in the simulation system consistent with the fluctuation dissipation theorem.

**Non-bonded interactions**

Throughout this section, we provide interaction potentials; in all cases, the forces associated with a given potential are determined by taking the gradient of the potential; the force between two objects is directed along the vector connecting their centers (or their surfaces in the case of membrane interactions).

*Fusion site clearing forces:* In order to prevent Syt molecules from blocking Syt-PM contact following disassembly of the Syt ring, constant repulsive forces push them away from the point of closest approach between the vesicle and the PM. We apply an interaction potential to Syt beads and the synaptic vesicle of the from

$$U_{\text{clearing}} = \begin{cases} F_0\,(\rho_0 - \rho), & \rho < \rho_0 \\ 0, & \rho \geq \rho_0 \end{cases}$$

where $\rho$ is the distance between the center of the vesicle the bead center projected onto the $XY$-plane and $F_0 = 0.05\;kT/\text{nm}$, and $\rho_0 = 10$ nm, approximately the radius of the SNARE ring. In simulations omitting this force, Syt molecules were distributed between the vesicle and PM, blocking vesicle-PM contact and inhibiting fusion.

*Excluded-volume interactions:* Forces prevent protein beads from overlapping with each other, the vesicle, and the plasma membrane.

For interactions between protein beads, the interaction potential is given by

$$U_{\text{exc}} = \begin{cases} \epsilon\left[\exp\left(-\dfrac{r^2}{2\sigma^2}\right) - \exp\left(-\dfrac{(1.15)^2}{2}\right)\right], & r < 1.15\,\sigma \\ 0, & r \geq 1.15\,\sigma \end{cases}$$

where $\epsilon = 10\;kT$ determines the strength of the interaction, $\sigma$ is the width of the interaction potential (given by the sum of the radii of the two beads), and $r$ is the distance between the centers of the beads. The parameter $\epsilon$ and the cutoff distance $1.15\sigma$ are chosen so that a fluctuation of magnitude $\sim 1\ kT$ will only lead to a small ~10% overlap of the beads.

For interactions between protein beads and membranes, the interaction potential is given by

$$U_{\text{lin}} = \begin{cases} \epsilon \dfrac{r - R_{\text{bead}}}{R_{\text{bead}}}, & r < R_{\text{bead}} \\ 0, & r \geq R_{\text{bead}} \end{cases}$$

where $r$ is the distance from the center of the bead to the membrane surface, $R_{\text{bead}}$ is the bead radius, and $\epsilon = 10\ kT$ is chosen so that a typical fluctuation of magnitude $\sim 1\ kT$ only leads to ~10% overlap.

*Electrostatic interactions:* The membranes as well as most protein beads are endowed with electrical charge. Due to the significant salt concentration inside of cells, charged elements in the simulation interact with each other via screened Coulomb forces.

The electrostatic interaction between the plasma-membrane and the vesicle is given by

$$U_{\text{el}} = R_{\text{ves}} Z_{\text{mb}} e^{-h/\lambda_D}$$

where $h$ is the separation between the vesicle and PM at the point of closest approach and $\lambda_D = 0.8$ nm is the Debye screening length for the monovalent physiological salt concentration, $[\text{salt}] = 0.15$ M (Leckband and Israelachvili, 2001). $Z_{\text{mb}}$ is calculated from the surface potential of the membranes $\psi_0$,

$$Z_{\text{mb}} = 9.38 \times 10^{-11} \tanh^2\left(\frac{\psi_0}{107}\right)\ \text{J/m},$$

where the surface potential is determined using the Graham equation $\psi_0 = 53.4 \sinh^{-1}\left(\frac{\sigma_{\text{m}}}{0.116[\text{salt}]^{1/2}}\right)$, and $\sigma_{\text{m}}$is surface charge density in $\text{C/m}^2$ (Israelachvili, 2011), determined using composition-weighted averages of the charge of the constituent lipids, Table S2.

The electrostatic potential between two CG protein beads is given by

$$U_{\mathrm{b,b}} = \frac{r_{\mathrm{bead}}}{2} Z^{i,j} e^{-d/\lambda_D},$$

where $d$ is the distance between the *surfaces* of the beads, and the parameter $Z^{i,j}$ is taken to be the geometric mean $Z^{i,j} = \pm\sqrt{\left|Z_{\mathrm{bead}}^{i} Z_{\mathrm{bead}}^{j}\right|}$, with the positive sign chosen for beads of like charge, and the negative sign chosen for beads of opposite charge (Israelachvili, 2011). The parameters $Z_{\mathrm{bead}}^{i}$ for individual protein beads are determined using a procedure identical to the one use for $Z_{\mathrm{mb}}$.

Lastly, the electrostatic potential energy between the vesicle membrane and a protein bead is given by

$$U_{b,\mathrm{ves}} = \frac{r_{\mathrm{bead}} R_{\mathrm{ves}}}{R_{\mathrm{ves}} + r_{\mathrm{bead}}} Z_{\mathrm{mb},b} e^{-d/\lambda_D},$$

where $d$ is the distance between the bead surface and the vesicle surface, and $R_{\mathrm{ves}} = 20$ nm is the radius of the vesicle. The parameter $Z_{\mathrm{mb},b}$ is calculated by taking the geometric mean $Z_{\mathrm{mb},b} = \pm\sqrt{\left|Z_{\mathrm{bead}}^{i} Z_{\mathrm{mb}}\right|}$, with the positive sign again chosen when the vesicle and bead have like charges, and the negative sign when they have opposite charges. Similarly, the electrostatic potential energy between a protein bead and the plasma membrane is given by

$$U_{b,\mathrm{PM}} = r_{\mathrm{bead}} Z_{\mathrm{mb},b} e^{-d/\lambda_D}.$$

*Van der Waals interactions:* The van der Waals interaction between the plasma-membrane and the vesicle is given by (Israelachvili, 2011)

$$E_{\mathrm{vdw}} = \frac{-A R_{\mathrm{ves}}}{6h},$$

where $A \approx 8 \times 10^{-21} J$ is the Hamaker constant for lipid bilayers and $R_{\mathrm{ves}} = 20$ nm is the radius of the vesicle. Again, $h$ is the separation between the PM and vesicle at the point of closest approach.

*Hydration layer-mediated interactions:* To bring the PM and the vesicle close together, a layer of water molecules that associate with the polar lipid head groups must be displaced. The hydration energy for vesicle-planar membrane is calculated by (Israelachvili, 2011)

$$U_{\mathrm{hyd}} = 2\pi\lambda_{\mathrm{hyd}}^{2} R_{\mathrm{ves}} P_0 e^{-h/\lambda_{\mathrm{hyd}}}$$

where, $\lambda_{\mathrm{hyd}} \approx 0.16$ nm, and $P_0 \approx 4.4 \times 10^4 \frac{kT}{\mathrm{nm}^3}$. The values of $P_0$ and $\lambda_{\mathrm{hyd}}$ are calculated using composition-weighted averages of the experimentally determined values of the constituent lipids in the vesicle and plasma membrane, Table S2 (Rand and Parsegian, 1989).

**Bond formation and bonded interactions**

Our simulations implement formation and dissociation of bonds between (1) SNARE complexes and Syt monomers, (2) Syt monomers and the PM, and (3) pairs of Syt monomers. Bonds are formed based on a distance criterion: when the beads making up the bond interface are within a predefined capture distance, a bond is created (Table S1). Bonded proteins are connected to each other by an elastic network of springs with rest length determined by the native state of the interaction (discussed in more detail below) (Wang et al., 2014; Zhou et al., 2015). Each bond then dissociates stochastically at a rate chosen to reproduce experimental measurements of the strength of the interaction.

In order to reproduce the experimentally measured dissociation constants, we performed simulations with one of each binding partner in a simulation box of fixed finite volume. The volume of the simulation box was chosen so that the concentration of binding partners is equal to the dissociation constant of the interaction. For the Syt-SNARE primary interaction, this volume corresponds to cubic box of side-length 48 nm, while for the Syt-Syt interaction, it corresponds to a box with side-length 118 nm. We then scanned over values of dissociation rate, $k_{\mathrm{off}}$ to find the value at which the molecules were bound with probability 0.5, Fig. S2 and Table S1.

*Elastic network:* Bonded proteins are connected with a collection of Hookean springs. The springs connect all beads within 1.5 nm of each other in the native state of the protein complex, with rest length equal to the distance between the beads in the native state. While the structure of the Syt-

Syt and Syt-SNARE complexes have been determined experimentally (Wang et al., 2014; Zhou et al., 2015), the flexibility of these bonds has not been measured to our knowledge. For Syt-membrane and Syt-SNARE bonds, we used the stiffest spring constants achievable in our simulations, Table S1; increasing the stiffness of the spring without decreasing the time step would cause un-physical oscillations.

However, the experimentally determined persistence length of Syt rings allows us to determine the spring constant for the Syt-Syt elastic network systematically. Electron micrographs of populations of Syt rings show a broad distribution of ring sizes (Wang et al., 2014; Wang et al., 2017; Zhu et al., 2021). The width of the ring size distribution can be used to infer the persistence length of Syt rings (Zhu et al., 2021). Experimentally observed distributions of Syt ring-sizes are consistent with a persistence length of $\sim 40 - 170$ nm; we therefore used a persistence length of $\sim$70 nm in our simulations, near the middle of the range of experimental values. Simulated rings of greater or lesser persistence length had no qualitative structural differences.

In order to impose the desired persistence length, we performed simulations of Syt rings, scanning over the Syt-Syt elastic network spring constant and measuring the resultant persistence length from fluctuations in the ring structure. Fluctuations in the angle $\theta$ between the tangent vector to the ring measured at adjacent monomers were then measured during simulations (n=5 runs per parameter value, $50$ μs per run). The persistence length of the Syt ring was inferred from the relation $L_\mathrm{p} = L_0/(\langle\theta^2\rangle - \langle\theta\rangle^2)$, where $\theta$ is the relative angle between tangent vectors at adjacent Syt monomers in the ring, $L_0$ is the distance between adjacent monomers (measured from their respective centers), and $\langle\cdot\rangle$ represents the expectation value. The spring constant was chosen to produce a persistence length of 70 nm, near the middle of the range of experimentally observed values, 40-170 nm (Wang et al., 2017; Zhu et al., 2021), Fig. S3 and Table S1.

**Determination of the persistence length from Syt ring fluctuations**

Consider a polymer of length $L$ and persistence length $l_p$. The bending energy of the polymer is given by

$$U = kT\, l_p \int_0^L \frac{\kappa^2}{2} \mathrm{d}s,$$

where $s$ is the arclength along the contour of the polymer, and $\kappa$ is the local curvature of the contour. Discretizing the polymer into segments of length $\delta l$, the bending energy is given by

$$U = kT\frac{l_p}{\delta l}\sum_i (1 - \cos(\theta_i)),$$

where $\theta_i$ is the angle between the unit tangent vectors at segment $i$ and segment $i + 1$. Taking $\theta_i \ll 1$, the energy is approximately

$$U \approx kT\frac{l_p}{\delta l}\sum_i \frac{\theta_i^2}{2}.$$

Thus, it follows by the equipartition theorem that

$$\langle\theta_i^2\rangle = \frac{\delta l}{l_p}.$$

Here, we used this formula to determine the persistence length of simulated Syt rings from fluctuations of the tangent vector, with $\delta l \approx 5$ nm corresponding to the distance between centers of the C2AB domains, Fig. S3. Adjusting the stiffness of elastic network connections allowed the persistence length to be tuned to within the experimental range.

**SNARE zippering kinetics**

At each time step, each SNARE complex can zipper or unzip one layer. Unzipping a layer removes the appropriate beads from the SNARE complex, and increases the contour length of the associated worm-like chain(s) representing the VAMP and Syntaxin LDs, while zippering causes the inverse. Which beads are added or removed from the SNARE complex depends on the layer which is zippering or unzipping. At the C-terminal end of the SNARE motifs (layers +4 through +8), unzipping a layer removes the VAMP, Syntaxin, and SNAP-25 beads from that layer, consistent with optical tweezers experiments showing that the C-terminal domain of the SNARE complex frays upon unzippering (Ma et al., 2015; Zhang et al., 2016). Unzipping other layers only removes VAMP beads.

In our simulations, zippering and unzipping of the SNARE complex is governed by a kinetic zipper model (Thompson et al., 1997). The rates of zippering and unzipping are

$$k_{\text{zip}} = k_0 e^{-\Delta E_{\text{zip}}/kT}$$

$$k_{\text{unzip}} = k_0$$

where $k_0 = 10^6\ \text{s}^{-1}$ is a typical time scale for assembly and disassembly of alpha helices (Gao et al., 2012; Kubelka et al., 2004), and the zippering energy $\Delta E_{\text{zip}}$ is the energy gained by assembling one additional layer onto the SNARE complex. The change in energy when zippering an additional bead is given by

$$\Delta E_{\text{zip}} = \Delta E_{\text{bind}} + \Delta E_{\text{linker}}$$

where $\Delta E_{\text{bind}}$ is the binding energy according to the zippering energy landscape determined by Ma et al. (Ma et al., 2015), and $\Delta E_{\text{linker}}$ is the change in the worm-like chain energy of the uncomplexed segment of VAMP. $\Delta E_{\text{bind}}$ was calculated using the number of uncomplexed VAMP residues as in previous studies (McDargh et al., 2018; Mostafavi et al., 2017).

**Syt $Ca^{2+}$-binding kinetics**

In our simulations, Syt monomers bind $Ca^{2+}$ ions at each of their five $Ca^{2+}$-binding sites, three of which are on the C2A domain, and two of which are on the C2B domain. All binding sites are assumed to have the same on-rate $k_{\text{on}}$; the two $Ca^{2+}$-binding sites on the C2B domain have the same off-rate, while the three sites on the C2A domain all have different off-rates, Table S1, as observed in many experimental studies (Arac et al., 2006; Millet et al., 2002; Radhakrishnan et al., 2009; van den Bogaart et al., 2012).

Experimentally reported values of the $Ca^{2+}$-binding affinity of the Syt C2B domain vary widely, and changes in the presence of different negatively charged lipids (Radhakrishnan et al., 2009; van den Bogaart et al., 2012). The on-rate of $Ca^{2+}$-binding has been measured to be ~$10^7 - 10^8\ \text{M}^{-1}\text{s}^{-1}$ (Chapman, 2008; Davis et al., 1999; Millet et al., 2002), while kinetic models of release at the calyx of Held suggest an on-rate of $\sim 2 - 5 \times 10^8\ \text{M}^{-1}\text{s}^{-1}$ (Bollmann et al., 2000; Lou et al., 2005; Schneggenburger and Neher, 2000; Wang et al., 2008). We simulated action potentials at the calyx of Held, scanning over these parameters to find the values that best reproduced the experimental probability of release $\sim 0.05 - 0.1$ and delay time $\sim 500$ μs (Lou et al., 2005; Schneggenburger and Neher, 2000, 2005; Wang et al., 2008). We found an on-rate of $k_{\text{on}} = 2 \times 10^8\ \text{M}^{-1}\text{s}^{-1}$ and a dissociation constant of $K_{\text{D}} = 25$ μM reproduced these observables well, both in the range of proposed values, Fig. S2 and Table S1.

**Prediction of EPSCs from simulated release distributions.**

EPSCs were predicted from simulations by determining the distribution of NT release times, and convolving the result with experimentally measured miniature EPSCs (mEPSCs). Distributions were determined from histograms in which release events were placed in 500 μs bins. Confidence intervals and SDs for EPSC amplitudes and delay times were determined using bootstrap re-sampling, with each distribution of release events resampled 200 times. Simulations in which release did not occur during the simulated time frame were included in the bootstrap resampling.

For simulations of $Ca^{2+}$ uncaging and AP-evoked release at the calyx of Held, we used the mEPSC measured in ref. (Schneggenburger and Neher, 2000), while for simulations of AP-evoked release from cerebellar granule cells, we used the mEPSC characterized in ref. (Malagon et al., 2016). We note that the mEPSC measured at the calyx of Held may have an unusually long tail, because these cells were treated with cyclothiazide in order to prevent desensitization of AMPA receptors, which is important for deconvolution analysis when large quantities of NT are released.

### SI Figures

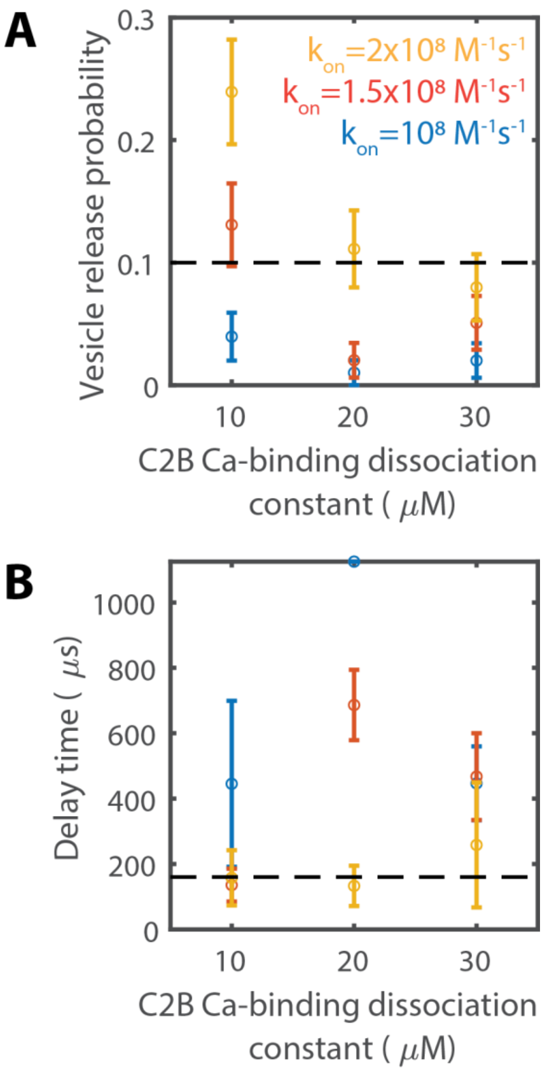


**Figure S1: Tuning Syt-Ca$^{2+}$ binding kinetics to reproduce release probability and delay time in the calyx of Held**

Vesicle release probability (A) and synaptic delay time (B) in simulations of AP-evoked release at the calyx of Held as a function of the Syt-Ca$^{2+}$ binding parameters $k_{\mathrm{on}}$ and $K_{\mathrm{D}} = k_{\mathrm{off}}/k_{\mathrm{on}}$ (n=100 per pair of parameter values). An on-rate of $k_{\mathrm{on}} = 2 \times 10^{8}\ \mathrm{M}^{-1}\mathrm{s}^{-1}$ was required to achieve the short delay times observed experimentally, and a dissociation constant $K_{\mathrm{D}} = 20\ \mu\mathrm{M}$ best reproduced the experimental vesicle release probability.

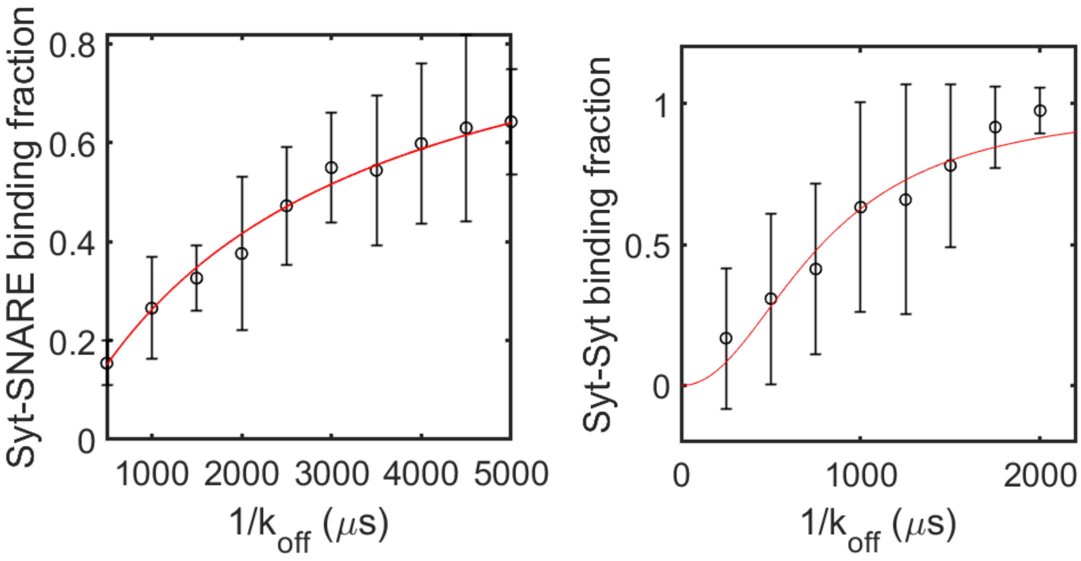


**Figure S2: Syt-Syt and Syt-SNARE binding kinetics are chosen to reproduce experimentally observed dissociation constants**

(A) Simulations of a single SNARE complex and a single Syt monomer in a finite simulation box with volume equal to the inverse of the experimentally observed dissociation constant, $K_{\mathrm{D}} = 14.8$ μM (Zhou et al., 2017); at this concentration, it is expected that the binding partners will be bound with probability 0.5. Syt-SNARE binding was diffusion controlled, i.e. a bond was formed when the beads comprising the primary interaction interface on the two binding partners were within a certain capture distance; dissociation occurred stochastically at a rate $k_{\mathrm{off}}$ that was varied in simulations, and the probability of binding was measured by dividing the total time the partners were bound by the total simulation time (n=10 runs per parameter value, 10 ms simulation time per run). The red curve represents a Hill function fit to the binding probability. The dissociation time that gave a binding probability of 0.5 was then chosen.

(B) Simulations of a pair of Syt monomers in a finite simulation box with volume equal to the inverse of the experimentally measured dissociation constant (Wang et al., 2017), varying the stochastic Syt-Syt dissociation rate $k_{\mathrm{off}}$ (n=10 runs per parameter value, 10 ms simulation time per run). The red curve represents a Hill function fit to the binding probability, with Hill coefficient $n = 2$. The dissociation time that gave a binding probability of 0.5 was selected.

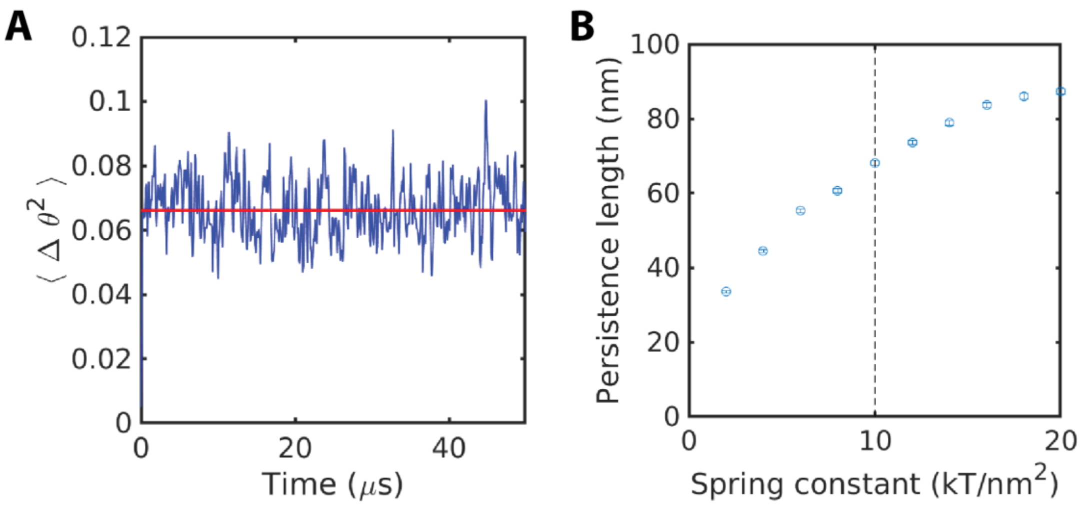


**Figure S3: Persistence length of Syt ring**

(A) Variance of the angle between the tangent vector to the Syt ring measured at adjacent Syt monomers as a function of time.

(B) Persistence length determined from simulations vs. elastic network spring constant. A spring constant of 10 kT/nm$^2$ produced the desired persistence length.

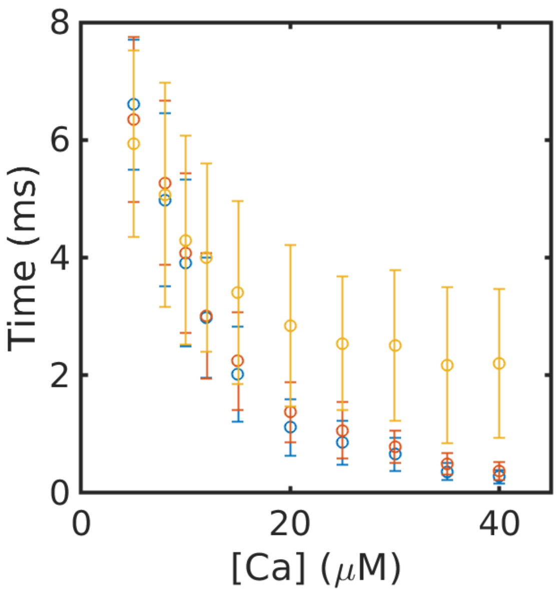


**Figure S4: Unclamping, assembly, and release time**

Average unclamping time (blue), SNARE-ring assembly time (orange) and fusion time (yellow) as a function of [$Ca^{2+}$]. The unclamping step grows significantly shorter as [$Ca^{2+}$] increases, while the time from unclamping to assembly remains ~0.2 ms independent of [$Ca^{2+}$], suggesting that the kinetics of the SNARE ring assembly process are insensitive to [$Ca^{2+}$].

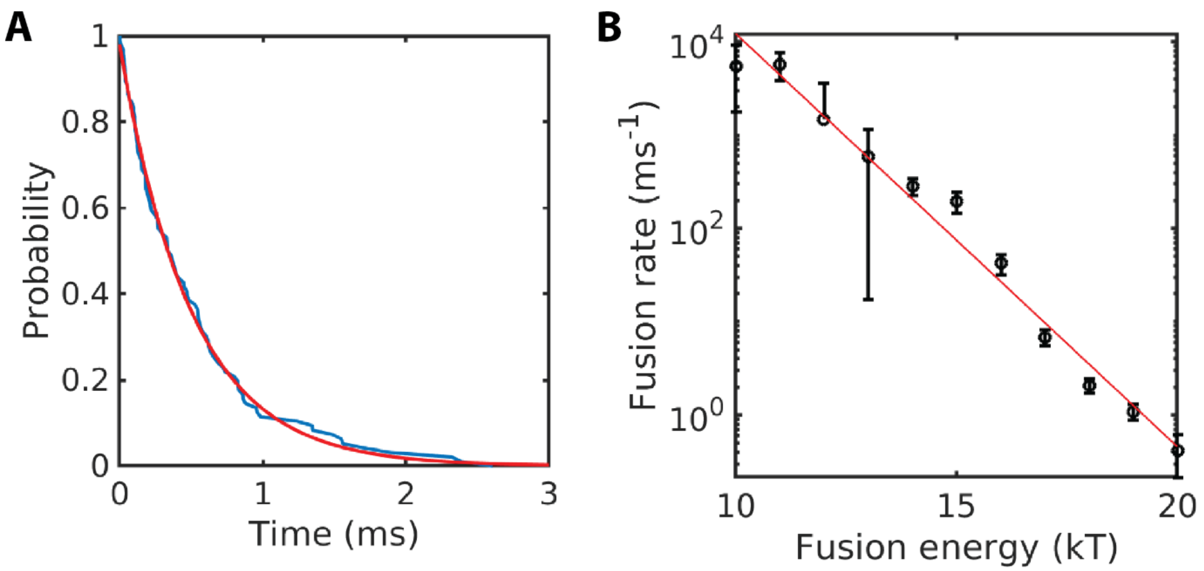


**Figure S5: SNARE-mediated fusion rates in simulations omitting Syt**

(A) Distribution of the length of the time interval between fusion events with a fusion energy of $20\ kT$ (blue) in a simulation without Syt and exponential fit of the distribution (red).

(B) Average fusion rate in the simulations in A as a function of activation barrier to fusion (black) and exponential fit to this data (red). The fusion rate scaled as $\sim \exp\left[-E_{\text{fusion}}/kT\right]$. Thus, decreasing the fusion threshold from $20\ kT$ to $19\ kT$ accelerated the fusion reaction by a factor of 2.6.

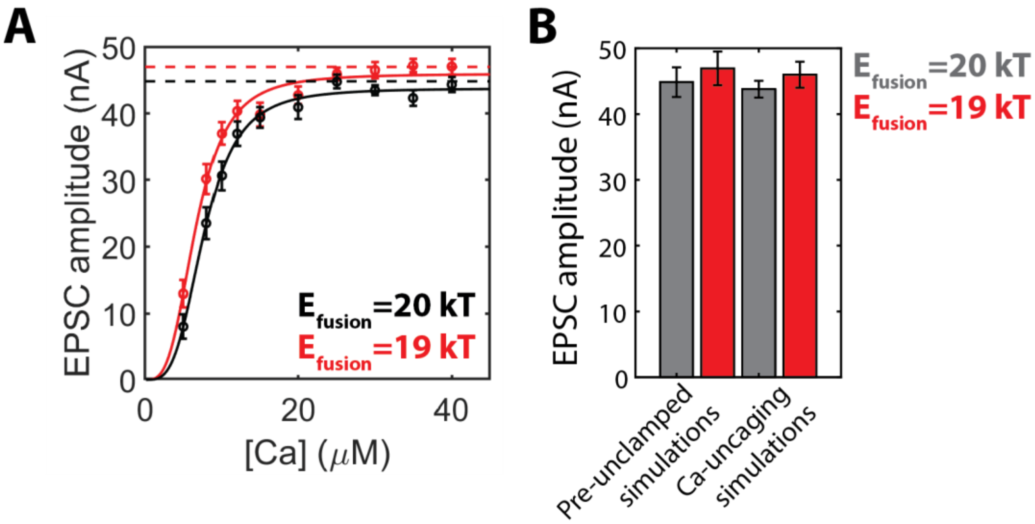


**Figure S6: SNARE-mediated membrane fusion is rate limiting at high [$Ca^{2+}$]**

(A) EPSC amplitude asymptotically approaches a fusion-limited value at high [$Ca^{2+}$]. EPSC amplitude as a function of [$Ca^{2+}$] in $Ca^{2+}$-uncaging simulations, with $E_{\text{fusion}} = 19\ kT$ (red) and $E_{\text{fusion}} = 20\ kT$ (black). Circular points represent measurements from simulations (n=100), while curves represent Hill function fits to the simulation data. Dotted lines show the EPSC amplitude measured from simulations in which all Syt $Ca^{2+}$-binding sites were assumed occupied at all times; these simulations were allowed to run 0.5 ms before data recording began; unclamping and SNARE ring assembly had therefore already completed at the beginning of data recording. Error bars represent 95% confidence intervals.

(B) Comparison of asymptotic EPSC amplitude determined from Hill function fits of $Ca^{2+}$-uncaging data with EPSC amplitude measured from simulations in which unclamping and SNARE ring assembly had completed prior to data recording. Error bars represent 95% confidence intervals.

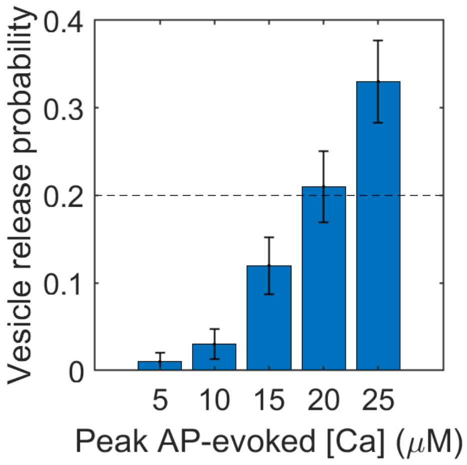


**Figure S7: The amplitude of the granule cell [$Ca^{2+}$] transient is tuned to reproduce the vesicle release probability**

Simulations of AP-evoked release at cerebellar granule cells were performed, approximating the [$Ca^{2+}$](t) time course as the experimentally observed $I_{\mathrm{Ca}}$ time-course(Kawaguchi and Sakaba, 2017). The normalization (i.e. the maximal [$Ca^{2+}$] value during the AP) was varied in simulations, and the vesicle release probability was measured ($n = 100$ simulations per parameter value). A peak [$Ca^{2+}$] of 20 μM reproduced the experimentally observed vesicle release probability.

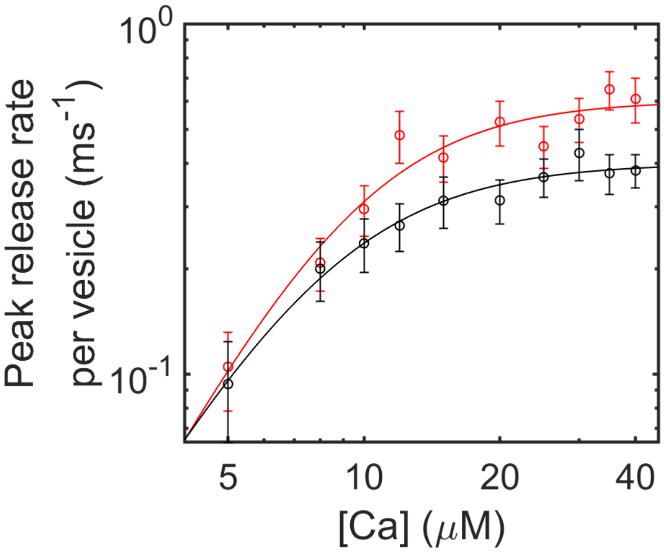


**Figure S8: Asymptotic vesicle release rate is increased by accelerating SNARE-mediated fusion**

Peak release rate as a function of [$Ca^{2+}$] in $Ca^{2+}$-uncaging simulations with $E_{\text{fusion}} = 20\ kT$ (black) and $E_{\text{fusion}} = 19\ kT$ (red), representing application of PDBu, with Hill function fits (smooth curves). Lowering the energetic barrier led to an increase in the asymptotic release rate at high [$Ca^{2+}$].

**Table S1: Model Parameters**

| **Parameter** | **Value** |
|---|---|
| $\delta t$, Time Step of MD simulation | 0.1 ns |
| $\eta$, Viscosity of water | $8.9 \times 10^{-3}$ dyn $\cdot$ s/cm$^2$ |
| $T$, Temperature | 298 K |
| $\lambda_\mathrm{D}$, Debye length[a] | 0.8 nm |
| $A$, Hamaker constant[b] | $8 \times 10^{-21}$ J |
| $l_\mathrm{p}$, Persistence length of worm-like chain[c] | 0.6 nm |
| Contour length per residue of uncomplexed regions[c] | 0.365 nm |
| $\lambda_\mathrm{hyd}$, Decay length for Hydration force[d] | 0.164 nm |
| $P_0$, Pressure prefactor for Hydration Force[d] | 44,273 $k_B T$/nm |
| Syt-SNARE capture distance | 0.7 nm |
| Syt-Syt capture distance | 1.5 nm |
| Syt-membrane capture distance | 0.8 nm |
| Syt-SNARE dissociation time[e] | 2800 μs |
| Syt-Syt dissociation time[e] | 1010 μs |
| C2A-membrane dissociation time[f] | 7.4 ms |
| C2B-membrane dissociation time[f] | 2.0 s |
| C2AB-membrane dissociation time[f] | 60 s |
| Synaptotagmin $Ca^{2+}$ binding rate, $k_\mathrm{on}$[g] | $2\times10^8$ $M^{-1}s^{-1}$ |
| $Ca^{2+}$ binding affinity of sites 1, 2, and 3 on C2A[h] | 120 μM, 465 μM, 1700 μM |
| $Ca^{2+}$ binding affinity of sites 1 and 2 on C2B[h] | 20 μM |
| Syt-PM capture distance | 0.8 nm |
| Syt-Syt capture distance | 1.6 nm |
| Syt-SNARE capture distance | 0.4 nm |

**Table S1.** (a) Calculated from physiological salt concentration of 0.15 M (Leckband and Israelachvili, 2001).

(b) Typical for lipid bilayers (Leckband and Israelachvili, 2001).

(c) Typical for unstructured polypeptides (Gao et al., 2012; Gebhardt et al., 2010).

(d) Determined from composition-weighted averages (Table S2) of values measured for pure lipid species (Rand and Parsegian, 1989).

(e) Chosen to reproduce experimental dissociation constants, Fig. S2.

(f) Off-rates measured in reference (Ma et al., 2017).

(g) Tuned to reproduce synaptic delay times at calyx of Held, Fig. S1; similar to on-rates in the literature (Bollmann et al., 2000; Davis et al., 1999; Lou et al., 2005; Schneggenburger and Neher, 2000).

(h) C2A values taken from ref. (Radhakrishnan et al., 2009); C2B values tuned to reproduce probability of release at calyx of Held; similar to affinities measured in the literature, Fig. S1 (Radhakrishnan et al., 2009; Rhee et al., 2005; van den Bogaart et al., 2012).

**Table S2: Membrane composition**

| | Vesicle | Plasma membrane |
|---|---|---|
| PE | 23% | 40% |
| PC | 36% | 40% |
| PS | 12% | 18% |
| PIP2 | 0 | 2% |

**Table S2.** Vesicle composition taken from (Takamori et al., 2006). Plasma membrane composition taken from (Hitzemann and Johnson, 1983).

#### Supplemental References

Arac, D., Chen, X., Khant, H.A., Ubach, J., Ludtke, S.J., Kikkawa, M., Johnson, A.E., Chiu, W., Sudhof, T.C., and Rizo, J. (2006). Close membrane-membrane proximity induced by Ca(2+)-dependent multivalent binding of synaptotagmin-1 to phospholipids. Nat Struct Mol Biol *13*, 209-217.

Bollmann, J.H., Sakmann, B., and Borst, J.G. (2000). Calcium sensitivity of glutamate release in a calyx-type terminal. Science *289*, 953-957.

Chapman, E.R. (2008). How does synaptotagmin trigger neurotransmitter release? Annu Rev Biochem *77*, 615-641.

Davis, A.F., Bai, J., Fasshauer, D., Wolowick, M.J., Lewis, J.L., and Chapman, E.R. (1999). Kinetics of synaptotagmin responses to Ca2+ and assembly with the core SNARE complex onto membranes. Neuron *24*, 363-376.

Gao, Y., Zorman, S., Gundersen, G., Xi, Z., Ma, L., Sirinakis, G., Rothman, J.E., and Zhang, Y. (2012). Single reconstituted neuronal SNARE complexes zipper in three distinct stages. Science *337*, 1340-1343.

Gebhardt, J.C., Bornschlogl, T., and Rief, M. (2010). Full distance-resolved folding energy landscape of one single protein molecule. Proc Natl Acad Sci U S A *107*, 2013-2018.

Hitzemann, R.J., and Johnson, D.A. (1983). Developmental changes in synaptic membrane lipid composition and fluidity. Neurochem Res *8*, 121-131.

Israelachvili, J.N. (2011). Intermolecular and surface forces: revised third edition (Academic press).

Kawaguchi, S.Y., and Sakaba, T. (2017). Fast Ca(2+) Buffer-Dependent Reliable but Plastic Transmission at Small CNS Synapses Revealed by Direct Bouton Recording. Cell Rep *21*, 3338-3345.

Kubelka, J., Hofrichter, J., and Eaton, W.A. (2004). The protein folding 'speed limit'. Curr Opin Struct Biol *14*, 76-88.

Leckband, D., and Israelachvili, J. (2001). Intermolecular forces in biology. Q Rev Biophys *34*, 105-267.

Lou, X., Scheuss, V., and Schneggenburger, R. (2005). Allosteric modulation of the presynaptic Ca2+ sensor for vesicle fusion. Nature *435*, 497-501.

Ma, L., Cai, Y., Li, Y., Jiao, J., Wu, Z., O'Shaughnessy, B., De Camilli, P., Karatekin, E., and Zhang, Y. (2017). Single-molecule force spectroscopy of protein-membrane interactions. Elife *6*.

Ma, L., Rebane, A.A., Yang, G., Xi, Z., Kang, Y., Gao, Y., and Zhang, Y. (2015). Munc18-1-regulated stage-wise SNARE assembly underlying synaptic exocytosis. Elife *4*.

Malagon, G., Miki, T., Llano, I., Neher, E., and Marty, A. (2016). Counting Vesicular Release Events Reveals Binomial Release Statistics at Single Glutamatergic Synapses. J Neurosci *36*, 4010-4025.

McDargh, Z.A., Polley, A., and O'Shaughnessy, B. (2018). SNARE-mediated membrane fusion is a two-stage process driven by entropic forces. FEBS Lett *592*, 3504-3515.

Millet, O., Bernado, P., Garcia, J., Rizo, J., and Pons, M. (2002). NMR measurement of the off rate from the first calcium-binding site of the synaptotagmin I C2A domain. FEBS Lett *516*, 93-96.

Mostafavi, H., Thiyagarajan, S., Stratton, B.S., Karatekin, E., Warner, J.M., Rothman, J.E., and O'Shaughnessy, B. (2017). Entropic forces drive self-organization and membrane fusion by SNARE proteins. Proc Natl Acad Sci U S A *114*, 5455-5460.

Radhakrishnan, A., Stein, A., Jahn, R., and Fasshauer, D. (2009). The Ca2+ affinity of synaptotagmin 1 is markedly increased by a specific interaction of its C2B domain with phosphatidylinositol 4,5-bisphosphate. J Biol Chem *284*, 25749-25760.

Rand, R.P., and Parsegian, V.A. (1989). Hydration Forces between Phospholipid-Bilayers. Biochimica Et Biophysica Acta *988*, 351-376.

Rhee, J.S., Li, L.Y., Shin, O.H., Rah, J.C., Rizo, J., Sudhof, T.C., and Rosenmund, C. (2005). Augmenting neurotransmitter release by enhancing the apparent Ca2+ affinity of synaptotagmin 1. Proc Natl Acad Sci U S A *102*, 18664-18669.

Schneggenburger, R., and Neher, E. (2000). Intracellular calcium dependence of transmitter release rates at a fast central synapse. Nature *406*, 889-893.

Schneggenburger, R., and Neher, E. (2005). Presynaptic calcium and control of vesicle fusion. Curr Opin Neurobiol *15*, 266-274.

Stein, A., Weber, G., Wahl, M.C., and Jahn, R. (2009). Helical extension of the neuronal SNARE complex into the membrane. Nature *460*, 525-U105.

Takamori, S., Holt, M., Stenius, K., Lemke, E.A., Gronborg, M., Riedel, D., Urlaub, H., Schenck, S., Brugger, B., Ringler, P., *et al.* (2006). Molecular anatomy of a trafficking organelle. Cell *127*, 831-846.

Thompson, P.A., Eaton, W.A., and Hofrichter, J. (1997). Laser temperature jump study of the helix<==>coil kinetics of an alanine peptide interpreted with a 'kinetic zipper' model. Biochemistry *36*, 9200-9210.

van den Bogaart, G., Meyenberg, K., Diederichsen, U., and Jahn, R. (2012). Phosphatidylinositol 4,5-bisphosphate increases Ca2+ affinity of synaptotagmin-1 by 40-fold. J Biol Chem *287*, 16447-16453.

Wang, J., Bello, O., Auclair, S.M., Wang, J., Coleman, J., Pincet, F., Krishnakumar, S.S., Sindelar, C.V., and Rothman, J.E. (2014). Calcium sensitive ring-like oligomers formed by synaptotagmin. Proc Natl Acad Sci U S A *111*, 13966-13971.

Wang, J., Li, F., Bello, O.D., Sindelar, C.V., Pincet, F., Krishnakumar, S.S., and Rothman, J.E. (2017). Circular oligomerization is an intrinsic property of synaptotagmin. Elife *6*.

Wang, L.Y., Neher, E., and Taschenberger, H. (2008). Synaptic vesicles in mature calyx of Held synapses sense higher nanodomain calcium concentrations during action potential-evoked glutamate release. J Neurosci *28*, 14450-14458.

Zhang, X., Rebane, A.A., Ma, L., Li, F., Jiao, J., Qu, H., Pincet, F., Rothman, J.E., and Zhang, Y. (2016). Stability, folding dynamics, and long-range conformational transition of the synaptic t-SNARE complex. Proc Natl Acad Sci U S A *113*, E8031-E8040.

Zhou, Q., Lai, Y., Bacaj, T., Zhao, M., Lyubimov, A.Y., Uervirojnangkoorn, M., Zeldin, O.B., Brewster, A.S., Sauter, N.K., Cohen, A.E., *et al.* (2015). Architecture of the synaptotagmin-SNARE machinery for neuronal exocytosis. Nature *525*, 62-67.

Zhou, Q., Zhou, P., Wang, A.L., Wu, D., Zhao, M., Sudhof, T.C., and Brunger, A.T. (2017). The primed SNARE-complexin-synaptotagmin complex for neuronal exocytosis. Nature *548*, 420-425.

Zhu, J., McDargh, Z.A., Li, F., Krishnakumar, S., Rothman, J.E., and O'Shaughnessy, B. (2021). Synaptotagmin rings as high sensitivity regulators of synaptic vesicle docking and fusion. bioRxiv, 2021.2003.2012.435193.
